# Differential Targeting of the Nucleosome Surface and Superhelical Crevice Sites with Ru and Os Organometallic Agents

**DOI:** 10.1101/2025.11.13.688318

**Authors:** Andrea Levy, Zenita Adhireksan, Thibaud von Erlach, Giulia Palermo, Alexey A. Nazarov, Christian G. Hartinger, Paul J. Dyson, Ursula Rothlisberger, Curtis A. Davey

**Affiliations:** Laboratory of Computational Chemistry and Biochemistry, Ecole Polytechnique Fédérale de Lausanne, Lausanne CH-1015, Switzerland; School of Biological Sciences, Nanyang Technological University, 60 Nanyang Drive, Singapore 637551; NTU Institute of Structural Biology, Nanyang Technological University, 59 Nanyang Drive, Singapore 636921; Institute of Chemical Sciences and Engineering, École Polytechnique Fédérale de Lausanne (EPFL), Lausanne, 1015 Switzerland; School of Chemical Sciences, University of Auckland, Private Bag 92019, Auckland 1142, New Zealand; Małopolska Centre of Biotechnology, Jagiellonian University, 31-007 Kraków, Poland

## Abstract

Platinum anticancer drugs tend to target DNA whereas certain ruthenium and osmium organometallic compounds, including those with known anticancer activity, preferentially bind histone proteins in chromatin. We earlier found that Ru/Os arene 2-pyridinecarbothioamide antitumor agents display unique or partially overlapping profiles of histone protein binding in the nucleosome compared to Ru arene phosphaadamantane (RAPTA) antimetastasis drugs, but the basis for this difference is unclear. Here we structurally characterized the nucleosome binding effects of arene ligand substitutions and carried out a multiscale simulation analysis, which reveals that the interplay between metal cation and non-leaving ligand identity dictates adduct stability and whether complexes target electronegative surface patches, internal crevices, or both. We show that the nucleosome superhelical crevice acts as a small molecule selectivity filter and that multi-site binding profiles can be expanded or reduced through defined ligand substitutions, which modulate dynamic and steric attributes. Our findings suggest new avenues for rationally developing Ru/Os organometallics that could help expand the scope of chromatin-targeting therapeutics.

**GRAPHICAL ABSTRACT:** 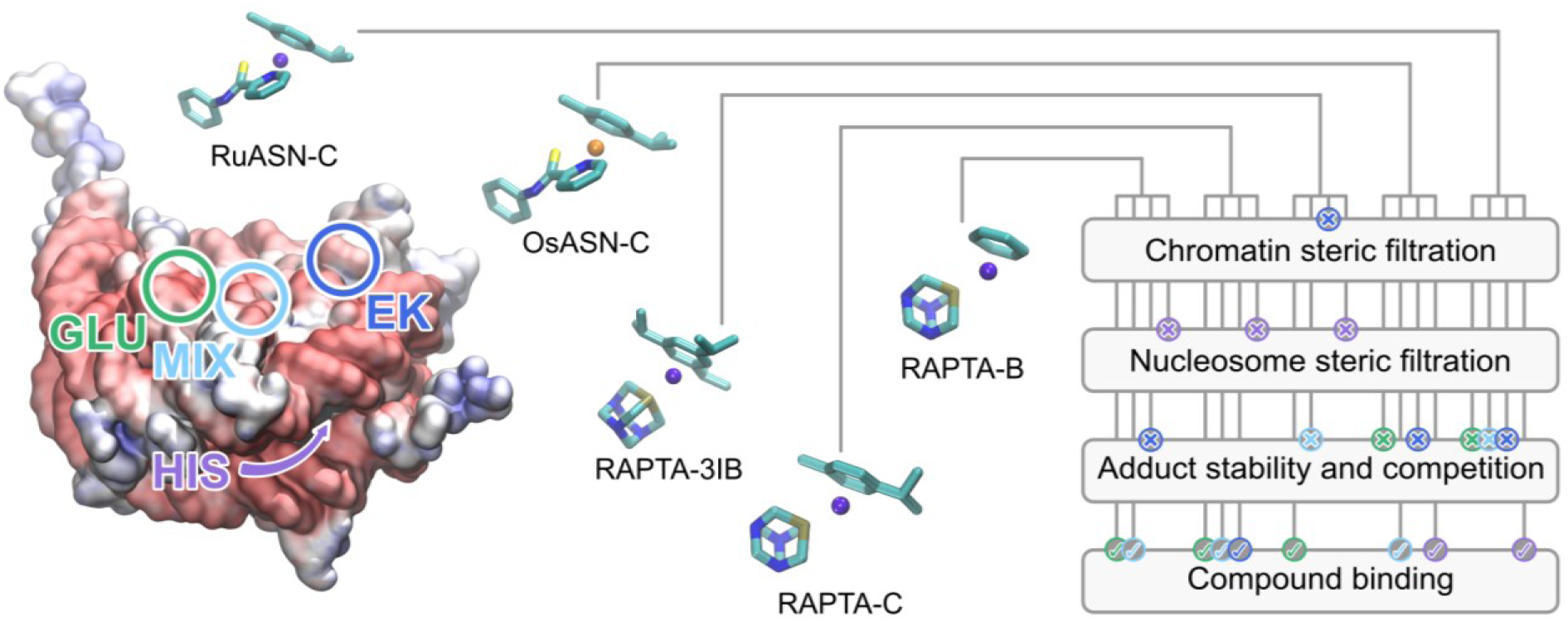

## INTRODUCTION

Metal-based drugs continue to attract considerable attention with many innovative non-classical approaches reported, and a number of excellent reviews are available (1-3). Nonetheless, platinum-based anticancer drugs, like cisplatin, oxaliplatin, and carboplatin, are among the most commonly used chemotherapeutic agents, which achieve efficacy by generating DNA cross-links (4,5). In spite of sustained progress in understanding the complex activity landscapes of the classic platinum drugs, these small molecules and analogous compounds form adducts in a relatively indiscriminate fashion and are limited by toxicity and resistance problems (6,7). Compounds based on alternative metal centres have generated interest because of their distinct mechanism of action compared to Pt drugs and selective activity against different cancers in combination with low toxicity; several agents having entered clinical trials (8-12). Given their unique, modular chemical/structural properties and biological activity profiles, ruthenium and osmium arene complexes have shown promise as a new class of chemotherapeutic agents (13-19).

Part of the intricacy associated with the activity profiles of genome-targeting agents arises from the packaging of DNA by histone proteins into chromatin. The repeating units of chromatin, nucleosomes, preside over a wealth of epigenetic information, which includes histone isoform/variant composition and a multitude of potential post-translational modifications that foster site-specific regulation of the genome (20,21). In fact, it has recently come to light that histone mutations and aberrant histone isoform/variant expression levels are common oncogenic drivers, implicating the histone proteins, or more specifically ‘oncohistones’, as bona fide cancer drug targets (22-25). Based on the success of Pt drugs, it was largely assumed that therapeutically active transition metal compounds would operate through targeting the DNA, but our initial studies on the interaction of Pt-, Ru-, Os-, and Au-based drugs/compounds with the nucleosome and chromatin revealed that certain agents have a preference for binding to the histone proteins as opposed to the DNA (6,26-34). Collectively, these investigations indicated a therapeutic potential in selective targeting of histones. Moreover, we found that the Pt and Au compounds investigated had distinct histone site preferences compared to each other as well as with respect to the set of Ru and Os arene complexes studied. The nucleosome site selectivity preferences for the Pt and Au agents could be attributed to the interplay of chemical predispositions of the compounds (i.e., type of metal and its associated oxidation state) with structural/dynamic (including allosteric) properties of the nucleosome (6,26,27,32,33). However, while we were able to rationalize why certain Ru/Os arene agents preferentially target the histones over the DNA (29-31,34), the basis of their histone site preferences remained unclear.

Here, we focus on two different types of Ru/Os arene compounds that notably display non-overlapping or partially overlapping histone site preferences depending on the exact nature of the agent. The RAPTA, Ruthenium(II)-Arene-PhosphaadamanTAne [(η^6^-*p*-arene)Ru(1,3,5-triaza-7-phosphaadamantane)Cl_2_], complexes (**Figure 1**) are antimetastasis drugs having inhibitory properties towards both primary and metastatic tumors as well as anti-angiogenic activity (13,35-37). RAPTA-C (C=cymene), RAPTA-T (T=toluene), and binuclear RAPTA agents bind selectively to electronegative histone sites on the nucleosome surface in vitro (**Figure 2**), while accumulating on the histone constituent of chromatin subsequent to uptake by cancer cells (28-30,32,38).

**Figure 1.**
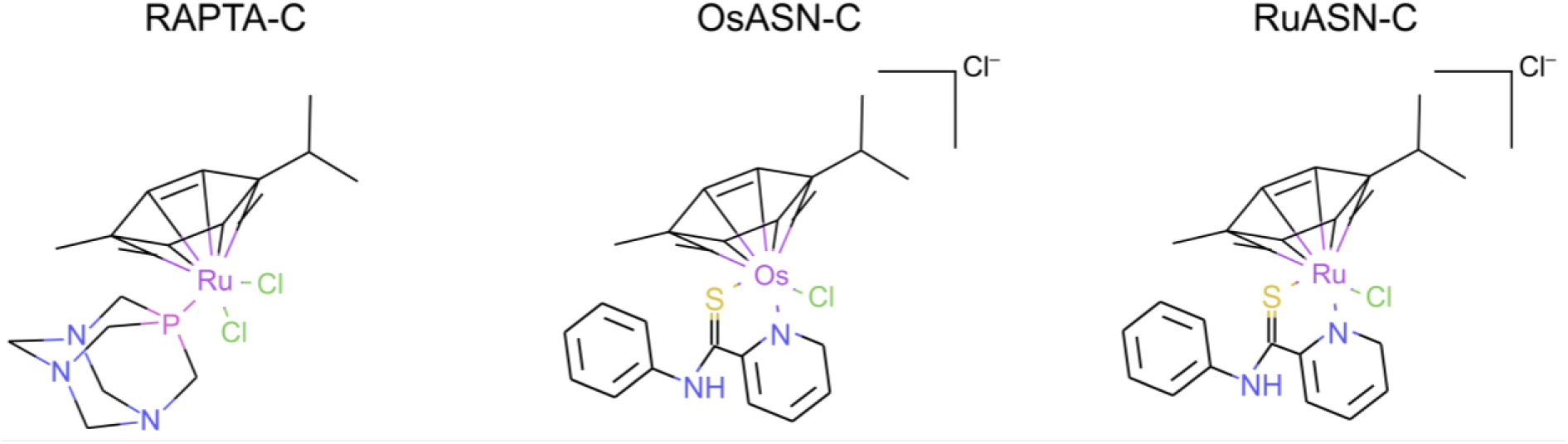
RAPTA and Ru/OsASN compounds. For the computational studies, the aquated version of the compounds has been used, with a total charge of +1 for the monoaquated RAPTA-C and +2 for the aquated Ru/OsASN-C compounds.

**Figure 2.**
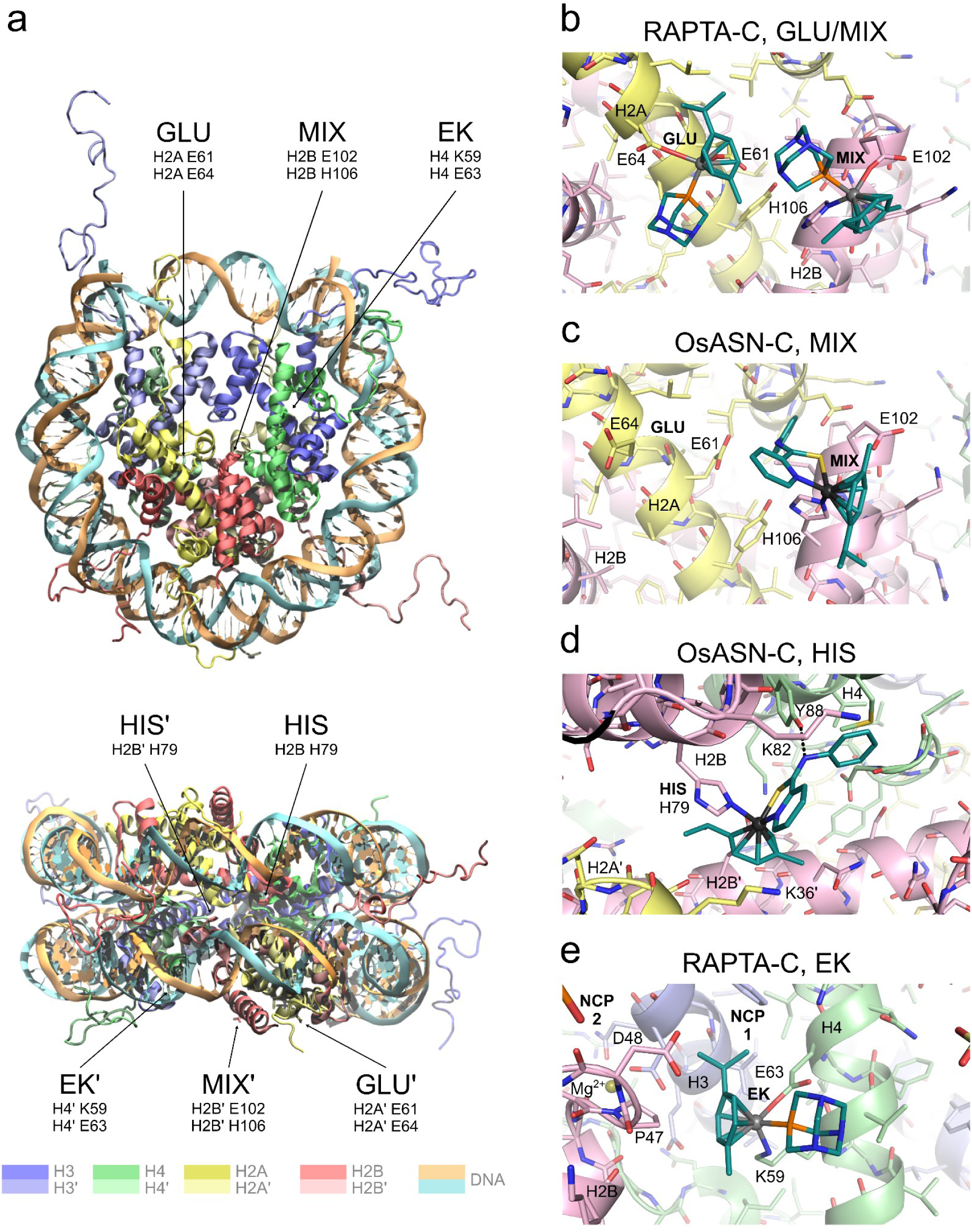
Nucleosome structure and organometallic compound-histone binding sites. **a)** Nucleosome core particle (NCP; PDB ID 1KX5; 39), highlighting the binding sites observed in crystal structures for Ru/Os arene compounds. GLU, MIX, EK, and their 2-fold symmetry equivalents (GLU’, MIX’, EK’) are surface sites, while HIS (and HIS’) are situated within the recessed nucleosomal crevice between the superhelical DNA gyres. **b–e)** NCP crystal structure-based details of RAPTA-C and OsASN-C (PDB ID 4J8U; 34) binding at the four different types of coordinating histone sites. The H3, H4, H2A, and H2B core histones are distinguished by pale colors of blue, green, yellow, and pink, respectively. A histone-non-leaving ligand hydrogen-bond is shown with a dashed line (d).

In contrast to RAPTA agents, RuASN-C [(η^6^-*p*-cymene)Ru(*N*-phenyl-2-pyridinecarbothioamide)Cl] and its osmium equivalent, OsASN-C [(η^6^-*p*-cymene)Os(*N*-phenyl-2-pyridinecarbo-thioamide)Cl] (**Figure 1**; A=arene, SN=sulfur/nitrogen bidentate ligand), display distinct cancer cell activity and localization attributes as well as histone binding profiles compared to the RAPTA agents (**Figure 2**; 34,40-42). While Ru-ASN-C and OsASN-C both display the intriguing ability to bind at internal (superhelical crevice) sites in the nucleosome, their histone binding and cancer cell activity profiles, as well as their resulting nucleosomal adduct structures, are not the same (34,41). This had prompted us to establish the basis for the differences both within and between these two Ru/Os arene compound types. In this work, we characterize the effect of non-leaving ligand substitutions by conducting crystallographic nucleosomal compound screening and carrying out a detailed multiscale analysis on the nucleosome-complex structures by employing molecular mechanical (MM), quantum mechanical (QM), and hybrid QM/MM techniques. The study reveals the basis for the site selectivity at specific carboxylate (glutamate) and imidazole (histidine) motifs, which can help rationalize the differential biological activities of the RAPTA, Ru-ASN, and Os-ASN agents and provides a guide for modulating their chromatin binding properties.

## MATERIALS AND METHODS

### Ruthenium compound synthesis

RAPTA agents having different arene ligands were synthesized as described previously (43). In brief, for the synthesis of RAPTA-X, where X is the arene group (either benzene [B], toluene [T], cymene [C], 1,3,5-trimethylbenzene [3MB], 1,3,5-triisopropylbenzene [3IB], or hexamethylbenzene [6MB]), equimolar amounts of 1,3,5-triaza-7-phosphatricyclo[3.3.1.1]decane and [Ru(η^6^-*p*-X)Cl_2_]_2_ were heated to reflux in methanol, and after cooling the product was precipitated by addition of diethyl ether.

### Crystallographic analysis of RAPTA-treated nucleosome

NCP crystals were produced and transferred into a stabilizing harvest buffer (37 mM MnCl_2_, 40 mM KCl, 20 mM K-cacodylate [pH 6.0], 24% [v/v] 2-methyl-2,4-pentanediol, and 2% [w/v] trehalose) as previously described (39,44). The 37 mM MnCl_2_ and 40 mM KCl buffer components were subsequently eliminated by gradual replacement with 10 mM MgSO_4_, followed by thorough rinsing of crystals with the MgSO_4_-containing buffer to remove any residual Mn^2+^ and Cl^−^ (28). The crystallographic analyses reported here stem from varied-duration incubations of NCP crystals with a 0.25–1.0 mM concentration of a given RAPTA compound included in the buffer (as outlined in **Tables 1 and 2**).

**Table 1.** Data collection and refinement statistics for NCP treated with RAPTA agents.

|  | **RAPTA-B** | **RAPTA-T** | **RAPTA-C** |
| --- | --- | --- | --- |
| **Data collection** |  |  |  |
| Space group | P2<sub>1</sub>2<sub>1</sub>2<sub>1</sub> | P2<sub>1</sub>2<sub>1</sub>2<sub>1</sub> | P2<sub>1</sub>2<sub>1</sub>2<sub>1</sub> |
| Cell dimensions |  |  |  |
| *a* (Å) | 106.68 | 106.68 | 106.35 |
| *b* (Å) | 110.01 | 109.82 | 109.86 |
| *c* (Å) | 182.43 | 182.34 | 182.59 |
| Resolution (Å) | 2.75–48.9<br>(2.75–2.90) | 2.60–60.8<br>(2.60–2.74) | 2.30–94.1<br>(2.30–2.42) |
| *R*<sub>merge</sub> (%) | 4.5 (124) | 6.1 (95.1) | 7.2 (103) |
| *R*<sub>pim</sub> (%) | 2.3 (78.3) | 2.8 (71.5) | 3.8 (89.5) |
| *I*/σ*I* | 20.1 (1.0) | 13.2 (1.0) | 8.6 (0.7) |
| Completeness (%) | 99.6 (98.0) | 88.4 (55.3) | 96.5 (80.2) |
| CC<sub>1/2</sub> (%) | 100 (44.7) | 99.9 (40.8) | 99.4 (57.6) |
| Redundancy | 6.0 (4.0) | 6.2 (3.1) | 4.8 (2.4) |
| **Refinement** |  |  |  |
| Resolution (Å) | 2.75–48.9 | 2.60–58.7 | 2.30–94.1 |
| No. reflections | 55,054 | 57,440 | 90,236 |
| *R*<sub>work</sub> / *R*<sub>free</sub> (%) | 22.0 / 26.9 | 21.9 / 24.2 | 22.7 / 25.6 |
| No. atoms | 12,075 | 12,077 | 12,126 |
| B-factors (Å<sup>2</sup>) | 115 | 102 | 91.7 |
| R.m.s. deviations |  |  |  |
| Bond lengths (Å) | 0.004 | 0.004 | 0.004 |
| Bond angles (°) | 1.38 | 1.31 | 1.41 |

Single crystal data sets. Values in parentheses are for the highest resolution shell.
**RAPTA-B:** 19-hour incubation of NCP crystal with a 1 mM concentration of RAPTA-B. **RAPTA-T:** 20-hour incubation of NCP crystal with a 0.25 mM concentration of RAPTA-T (32). **RAPTA-C:** 56-hour incubation of NCP crystal with a 0.75 mM concentration of RAPTA-C (28).

**Table 2.** Data collection and refinement statistics for NCP treated with RAPTA agents.

|  | **RAPTA-3MB** | **RAPTA-6MB** | **RAPTA-3IB** |
| --- | --- | --- | --- |
| **Data collection** |  |  |  |
| Space group | P2<sub>1</sub>2<sub>1</sub>2<sub>1</sub> | P2<sub>1</sub>2<sub>1</sub>2<sub>1</sub> | P2<sub>1</sub>2<sub>1</sub>2<sub>1</sub> |
| Cell dimensions |  |  |  |
| *a* (Å) | 106.86 | 107.11 | 106.78 |
| *b* (Å) | 109.91 | 110.03 | 109.90 |
| *c* (Å) | 182.44 | 182.33 | 182.34 |
| Resolution (Å) | 2.70–48.9<br>(2.70–2.85) | 2.99–76.7<br>(2.99–3.15) | 2.70–76.6<br>(2.70–2.85) |
| *R*<sub>merge</sub> (%) | 4.0 (115) | 6.4 (93.0) | 9.8 (77.2) |
| *R*<sub>pim</sub> (%) | 2.0 (66.9) | 3.3 (61.7) | 4.9 (54.6) |
| *I*/σ*I* | 24.5 (1.1) | 11.3 (1.2) | 8.6 (1.1) |
| Completeness (%) | 96.9 (80.6) | 99.3 (95.7) | 86.4 (47.9) |
| CC<sub>1/2</sub> (%) | 100 (60.9) | 99.7 (55.8) | 98.6 (49.2) |
| Redundancy | 6.0 (4.0) | 5.4 (3.3) | 5.2 (2.8) |
| **Refinement** |  |  |  |
| Resolution (Å) | 2.70–48.9 | 2.99–70.8 | 2.70–76.6 |
| No. reflections | 56,715 | 43,044 | 50,290 |
| *R*<sub>work</sub> / *R*<sub>free</sub> (%) | 21.8 / 27.2 | 21.8 / 26.8 | 23.0 / 29.1 |
| No. atoms | 12,081 | 12,064 | 12,067 |
| B-factors (Å<sup>2</sup>) | 110 | 122 | 102 |
| R.m.s. deviations |  |  |  |
| Bond lengths (Å) | 0.004 | 0.004 | 0.004 |
| Bond angles (°) | 1.40 | 1.46 | 1.35 |

Single crystal data sets. Values in parentheses are for the highest resolution shell.
**RAPTA-3MB:** 3-hour incubation of NCP crystal with a 1 mM concentration of RAPTA-3MB. **RAPTA-6MB:** 4-hour incubation of NCP crystal with a 1 mM concentration of RAPTA-6MB. **RAPTA-3IB:** 21-hour incubation of NCP crystal with a 1 mM concentration of RAPTA-3IB.

Single-crystal X-ray diffraction data were recorded as described previously (29), with crystals mounted directly into the cryocooling N_2_ gas stream set at −175 °C, using beam lines X06SA or X06DA of the Swiss Light Source (Paul Scherrer Institute, Villigen, Switzerland) and an X-ray wavelength of 1.5 Å. Data were processed with MOSFLM (45) and SCALA (46) from the CCP4 package (47,48). The native NCP models (PDB codes 2NZD and 3REH; 28,44) and NCP with bound RAPTA-C (PDB code 3MNN; 28) and RAPTA-T (PDB code 5DNM; 32) were used for initial structure solution by molecular replacement and comparative analysis. Structural refinement and model building were carried out with the COOT (49) and REFMAC (50) programs from the CCP4 suite (47,48).

Refinement restraint parameters for the RAPTA species were based on the small-molecule crystal structures (43,51) and included bond length restraints to the coordinating histone groups. The RAPTA-C and RAPTA-T structures described here differ from those reported earlier (28,32) by virtue of the imposition of bonding restraints to the coordinating glutamate, histidine, and/or lysine groups as well as extension of the diffraction data resolution. Graphic figures were prepared with PyMOL (DeLano Scientific LLC, San Carlos, CA, USA). Data collection and structure refinement statistics are given in **Tables 1** and **2**.

### Models for computational investigations

The system preparation and force field parameters were chosen similarly to previous studies on the same system (29,32,52). In brief, the apo system was initially obtained by refining the structure of the native NCP solved at 2.80 Å resolution (PDB code 1AOI; 53) and modeled with the AMBER force field ff14SB (54,55) with the ff99bsc0 modifications for DNA (56). The TIP3P model (57) was employed for the description of explicit waters, with the addition of Na^+^ counterions to neutralize the total charge, leading to a total number of more than ∼200,000 atoms. The parameters for the different drugs were adapted from previous studies (58). As aquation is a process happening spontaneously in the cell medium (59,60; activating, therefore, the prodrug), the simulations were initiated with a monoaquated, positively charged version of RAPTA-C. Similarly, experimental results have shown that the aquation of the OsASN-C compound is a thermodynamically favorable reaction, and also in this case, the aquated version of the compound has been simulated, using the R-enantiomer, which is the dominant adduct observed in the crystal structure.

Classical MD simulations were performed with GROMACS (61-63), heating the apo system to 300K (NVT ensemble), and then equilibrating for hundreds of ns in the NPT ensemble. Then, the different drugs were added to the bulk solvent away (∼40 Å) from the system and let equilibrate. The distance between the drug and the target location has then been constrained to perform thermodynamic integration (discussed in more detail in the next section) and decreased systematically. Every frame has been equilibrated, and the trajectory has been used to calculate the corresponding force due to the distance constraint. For each window, at least ∼10ns were simulated, or until the constraint forces converged to their average values.

Born–Oppenheimer (BO) QM/MM MD simulations were performed with CPMD (64) with the QM/MM electrostatic embedding scheme by Laio et al (65). Simulations were started from a configuration from a previous constrained MD run at the MM level. The QM level of choice was Density Functional Theory (DFT), using the BLYP functional (66,67) with norm-conserving Martins-Trouiller pseudopotentials (68) and correcting the exchange-correlation functional through the use of properly parameterized Dispersion Corrected Atom-Centered Potentials (69). The atoms treated at the QM level include the drug and the binding amino acid side chains, i.e., H2A E61 and H2A E64 for the GLU site, H2B H106 and H2B E102 for the MIX site, and H2B H79 for the HIS site. The bonds crossing the QM–MM boundary have been treated with monovalent pseudopotentials on the C_α_ atoms. Wavefunctions were expanded in a plane wave basis with a cutoff of 70Ry for RAPTA-C, and 100Ry for OsASN-C. The QM/MM simulations consisted of an initial wavefunction optimization, followed by thermalization of the system at 300 K. Then Born–Oppenheimer (BO) MD simulations were carried out with a timestep of 10*ħ*/*E_ℎ_* (∼0.24 fs). The QM and the MM subsystems were thermostated by three separate chains of Nosé–Hoover thermostats (70,71), for the QM atoms, the NCP, and the solvent, respectively. Analogously to the MM case, the distance between the drug and the target location has been successively constrained to perform thermodynamic integration. For each window, at least ∼1 ps were simulated, or until the constraint forces converged to their average values.

### Multilevel thermodynamic integration

Thermodynamic integration (TI) is a well-established method for free energy calculations from molecular dynamics simulations (72,73). In brief, the free energy difference along a reaction path defined by varying a reaction coordinate *ξ* can be obtained as

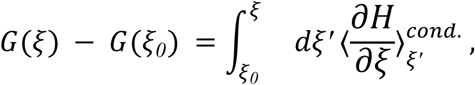

where 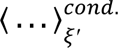 indicates a conditional average evaluated at *ξ*(*r*) = *ξ′*, which can be conveniently evaluated in the so-called blue-moon ensemble as the time average over a constrained trajectory, taking into account an additional factor to compensate for the bias introduced by the constraint. However, if the reaction coordinate is chosen as the distance between two atoms *i* and *j*, i.e. *ξ* = |*r*_*i*_ − *r*_*j*_|, the equations become simpler and the mean force can be directly estimated from the time average of the constraint force without the need for reweighting or correction terms.

TI within the blue moon ensemble has been extensively applied to MD simulations at MM (74), QM (75,76), and QM/MM level (77-79). The approach we propose in this work, *multilevel TI*, is a combination of an MM TI performed at large distances, where long sampling time is needed to ensure convergence of the constraint force, together with a QM/MM TI at short distances, where the MM description is not accurate enough and is not able to describe, e.g., bond breaking or formation. The total profile is obtained by combining the constraint forces before performing the integration to get the free energy, assuming that for large enough distances, the constraint forces from an MM-only simulation and a QM/MM one should match. This can be verified by always ensuring an overlapping region between the constrained distances studied at MM and QM/MM levels, enabling us to verify that the profile from the QM/MM forces is similar to or lower than the MM one in the overlap region.

The main advantage of multilevel TI is the possibility of getting a full free energy profile, from the bulk solvent to the binding site. Moreover, performing MM TI first allows us to get information about free energy barriers involved in accessing a putative binding site. If the free energy barrier from MM TI is already large, it is not helpful to proceed with a long and computationally costly QM/MM TI, since the binding site is unlikely to be reached in the first place. This is in line with the philosophy of the QM/MM approach of focusing the computational effort where it is strictly needed.

### Molecular volume and intermolecular contact analyses

For the molecular volume comparison between RAPTA-C and OsASN-C, we carried out a Bader’s atom in molecule analysis. For each compound, a representative conformation from the TI window at 40 Å was extracted, where the ligand is in the solvent. From the conformation, we generated a cube file containing the electron density generated using the same parameters as those in the subsequent QM/MM simulations and analysed them with the Bader Charge Analysis code (Version 1.05 08/19/23, available at https://theory.cm.utexas.edu/henkelman/code/bader/).

We also performed a contact analysis with an in-house Python code distributed in a Jupyter Notebook (deposited on Zenodo https://zenodo.org/records/17258752). The analysis consists of using a fixed structure as a reference, and conformations from the MD, and computing for each frame the distance for the closest contact, i.e. lowest distance among atoms of the reference structure vs MD, and the number of contacts, i.e. the number of atoms of the two structures with distances lower than a threshold (2.0 Å in this case). Before the contact analysis, we aligned the conformations from the MD to the crystal structure based on Cα carbons of the protein using VMD (80). Then we extracted the reference conformation of the drug at the MIX site and the conformations of the drug at the GLU site from the MD simulations.

### Binding energy QM calculations

To compute the influence of the Ru vs Os metal center at the HIS site, we computed the energy of binding for RuASN-C and OsASN-C compounds with QM calculations on a minimal QM model of the system, composed of the drug and the binding histidine. More details, together with the results for all reactants and products selected, are reported in the Supporting Information. Briefly, to compute the binding energy of the arene compound, we used as reactants the arene itself and the non-arene compound with Ru/Os bound to histidine, and as product Ru/OsASN-C bound to histidine. Similarly, for the binding energy of the non-arene compounds, we used the non-arene compound and the arene one bound to histidine as reactants, and Ru/OsASN-C bound to histidine as the product.

We compared results with different DFT functionals, namely BLYP (66,67), B3LYP (66,67,81,82), M06 (83), and M06L (84), while performing a full geometry optimization on the different reactants and products using a 6-31G* basis set and LANL2DZ pseudopotentials to describe the transition metal. The calculations have been performed using Gaussian 09 (85).

## RESULTS

### Ru/Os Arene Agent-Histone Binding Sites on the Nucleosome

We originally observed that RAPTA-C, RAPTA-T, and binuclear RAPTA compounds commonly coordinate to glutamate sites located in highly electronegative regions on the nucleosome (28-30,32,33), whereas RuASN-C and OsASN-C are seen to coordinate only to histidine (34; **Figures 1 and 2**). Two of the binding sites, referred to here as the GLU and the MIX sites, are situated in the nucleosome acidic patch, a key regulatory site on the surface of the H2A-H2B dimers rich in glutamate and aspartate residues (20,21; **Supplementary Figure S1**). The MIX site is common to the coordination of both classes of compounds, as it is composed of both a glutamate and a histidine side chain, rendering it capable of engaging in either monodentate or bidentate metal cation coordination (**Figures 2a–c**). Conversely, the other RAPTA-binding site (the GLU site) in the acidic patch is composed of two glutamate side chains and is capable of bidentate metal cation coordination. Distinct from the surface location of the acidic patch GLU and MIX sites, an internal motif (the HIS site) consisting of a single histidine side chain is present in the nucleosome superhelical crevice (**Figures 2a & 2d)**. A fourth type of histone coordinating site (the EK site) involves a dual carboxylate/amine motif, which is unique to the binding of RAPTA-C (**Figures 2a & 2e**). From the pseudo-two-fold symmetry of the nucleosome, the GLU, MIX, HIS, and EK sites each have a symmetry-related counterpart (GLU’, MIX’, HIS’, and EK’; **Figure 2a**).

### Nucleosome Crevice Acts as a Small Molecule Filter

Given the intriguing nature of site selectivity between the different classes of Ru/Os arene compounds, we sought to understand the factors that govern the binding selectivity. We first asked the question why RAPTA compounds do not bind to the HIS site. The symmetry-related HIS (H2B H79) and HISʹ (H2B′ H79) sites are ∼21 Å apart and are situated within a narrow crevice that is only accessible by diffusion through the two juxtaposed DNA gyres (**Figure 2a**). In order to assess the full free energy profile of compound binding from the bulk solvent to the binding sites, we conducted multilevel thermodynamic integration (TI) combining constrained molecular dynamics (MD) simulations at MM and QM/MM levels (additional details on multilevel TI are provided in the Materials and Methods section). The free energy profile, obtained using a classical (MM) description for RAPTA-C entry of the nucleosome crevice and approach towards the HIS site, revealed a large energy barrier associated with the penetration of the compound through the confined histone binding pocket (**Figure 3a and Supplementary Figure S2**). The free energy of approach peaks at a distance of around 12 Å from the HIS site, where steric clashes with the bulky non-leaving ligands of RAPTA-C are prevalent. An additional barrier associated with the RAPTA-C coordination reaction at H2B H79 would translate to a high overall energetic barrier for binding to the HIS site.

**Figure 3.**
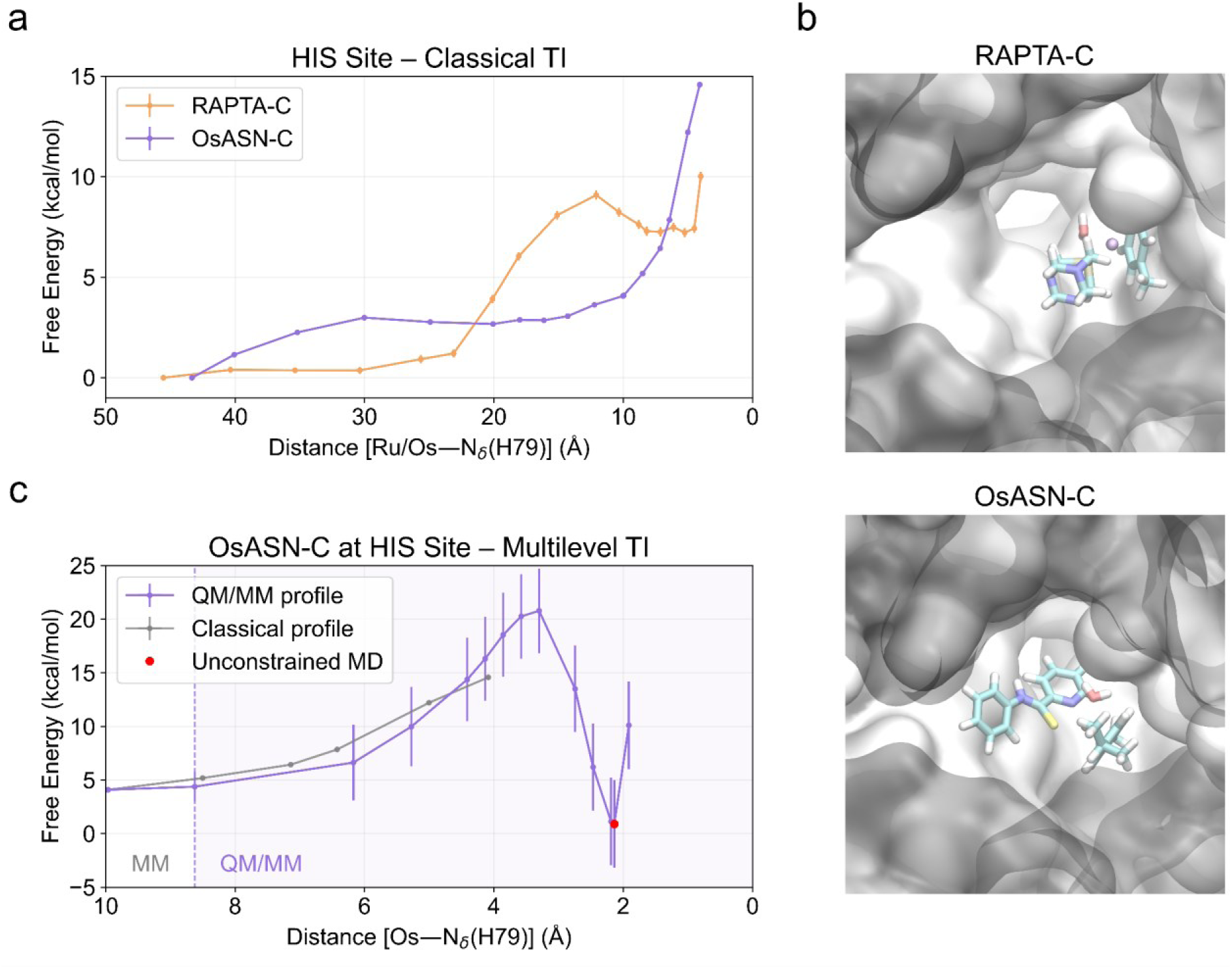
RAPTA-C and OsASN-C binding at the HIS site. **a)** Free energy profiles from classical TI. **b)** RAPTA-C and OsASN-C conformations at the entrance of the channel. Snapshot from classical TI. **c)** Multilevel free energy profile for OsASN-C, combining classical TI, QM/MM TI, and unconstrained QM/MM MD simulations. The region in which QM/MM TI is performed is highlighted with a shaded background.

Unlike the RAPTA compounds, OsASN-C and RuASN-C both form adducts at the HIS sites in crystallographic experiments. However, the resulting RuASN-C species at H2B/H2B′ H79 appear to have their bidentate SN ligands (*N*-phenyl-2-pyridinecarbothioamide or *N*-fluorophenyl-2-pyridinecarbothioamide) cleaved, in contrast to those of OsASN-C, which retain both the cymene and SN ligands (34). This is consistent with QM calculations performed with different DFT functionals, where we compute the binding energy of the arene (η^6^-*p*-cymene) and non-arene (*N*-phenyl-2-pyridinecarbothioamide) ligands in RuASN-C and OsASN-C (**Supplementary Table S1**). Notably, we observed a reduced SN ligand binding energy for RuASN-C relative to OsASN-C.

Since SN ligand cleavage in solution is expected to influence site selectivity, especially steric access, we performed MM TI calculations on OsASN-C to compare with the activity profile of RAPTA-C. In contrast to RAPTA-C, OsASN-C does not display a steric challenge, as it passes through the histone channel bottleneck with only a small increase in free energy (**Figure 3a**). Subsequent TI calculations employing a QM/MM treatment to capture the complete reaction profile (including coordination bond formation) reveal a modest overall free energy barrier of ∼20 kcal/mol associated with Os coordination at the H2B H79 δ-nitrogen atom and departure of the water ligand (**Figure 3c and Supplementary Figure S3**). The resulting OsASN-C adduct conformation remains stable when constraints are removed and is consistent with that of the crystal structure.

The TI analysis shows that the differential HIS site binding activity between the RAPTA and Ru/OsASN compounds arises from the presence of both arene and PTA non-leaving ligands, which create a roughly spherical and rigid steric profile. In this way, the nucleosome crevice, harboring a narrow access channel to the HIS site, acts as a steric site selectivity filter. Indeed, while OsASN-C has a similar overall steric volume as RAPTA-C (∼1.37×10^4^ Å^3^ from a Bader’s atoms in molecules analysis [86]; **Table 3**), it displays a significant degree of conformational freedom, which RAPTA-C does not. In fact, OsASN-C is able to adopt an altered non-leaving ligand configuration, rendering it with a flat structural profile that facilitates facile diffusion through the crevice bottleneck (**Figures 3b, 4, and Supplementary Video 1**).

**Table 3.** Results from Bader’s atom in molecule analysis for RAPTA-C and OsASN-C.

| Parameter | RAPTA-C | OsASN-C |
| --- | --- | --- |
| Number of atoms | 51 | 53 |
| Grid size | $241 \times 241 \times 241$ | $241 \times 241 \times 241$ |
| Number of Bader maxima | 221,123 | 240,632 |
| Significant maxima | 108 | 112 |
| Total number of electrons | 141.0000 | 151.0000 |
| Vacuum charge | 0.0000 | 0.0000 |
| Bader's volume ($\text{\AA}^3$) | $1.37 \times 10^4$ | $1.37 \times 10^4$ |

**Figure 4.**
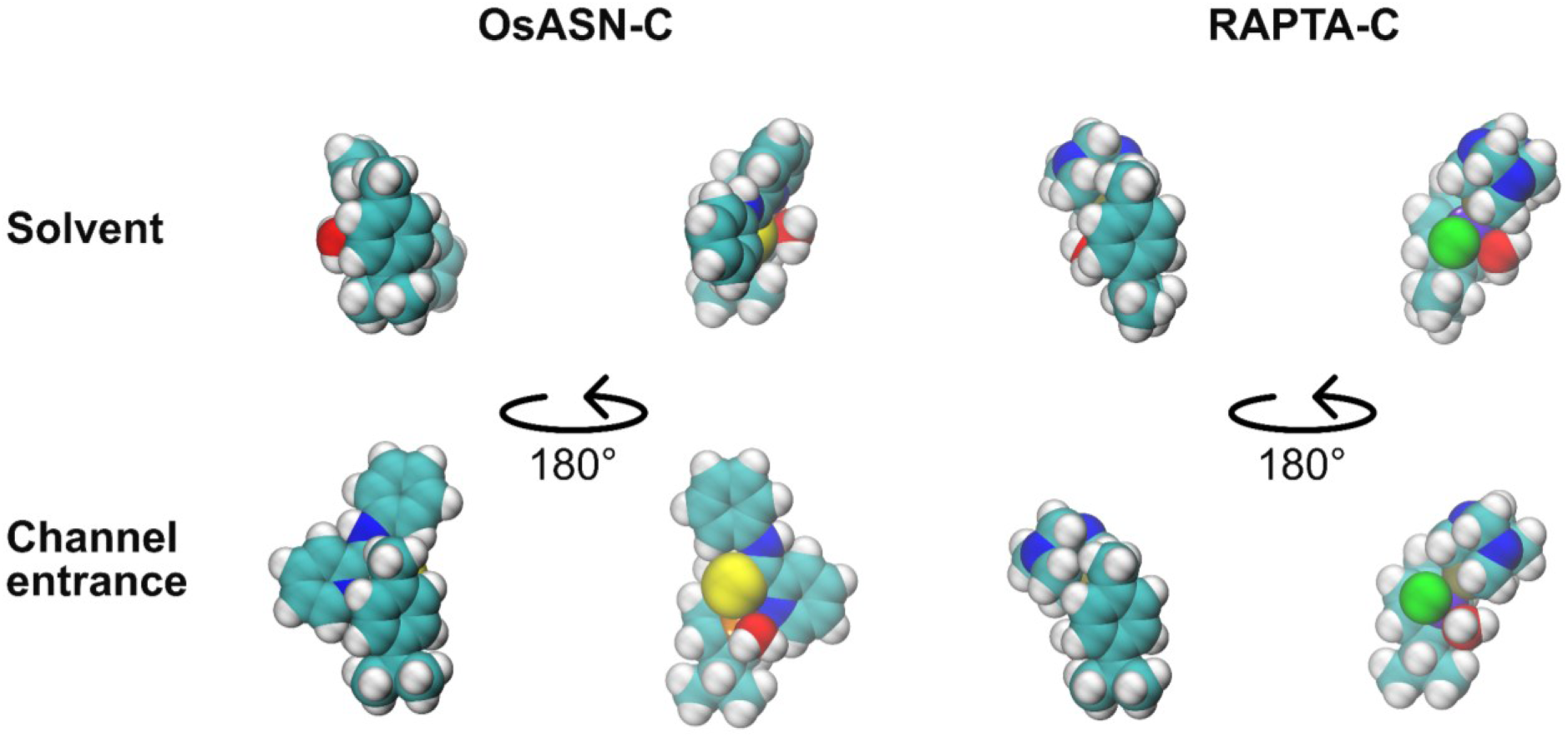
Comparison of compound conformations between the bulk solvent and at the channel entrance to the nucleosome superhelical crevice HIS site. The conformations have been extracted from TI windows at 40 Å (bulk solvent) and at 12 Å (channel entrance).

### Neighboring Sites Demand Mutual Compatibility

In contrast to the HIS sites, the GLU and MIX sites are freely accessible from the bulk solvent, confirmed by the barrier-free approach from MM-TI, of either RAPTA-C or OsASN-C up to a distance of around 6 Å from the binding sites (**Figures 5a-b and Supplementary Figure S4**). Following up with QM/MM, multilevel TI (**Figures 5c-d and Supplementary Figure S5**) reveals a small barrier for OsASN-C coordination at the H2A E61 carboxylate oxygen atom of the GLU site and an even lower barrier to its coordination at the H2B H106 ε nitrogen atom of the MIX site. The latter is associated with a deep energetic minimum (−15 kcal/mol) and a final adduct configuration compatible with the crystal structure. This rationalizes the proclivity of OsASN-C for binding at the MIX site, leaving the question as to why this agent is not seen to bind to the adjacent GLU site in the crystallographic analyses, even if, from the QM/MM TI profile, its binding seems possible but less strong than the one at the MIX site.

**Figure 5.**
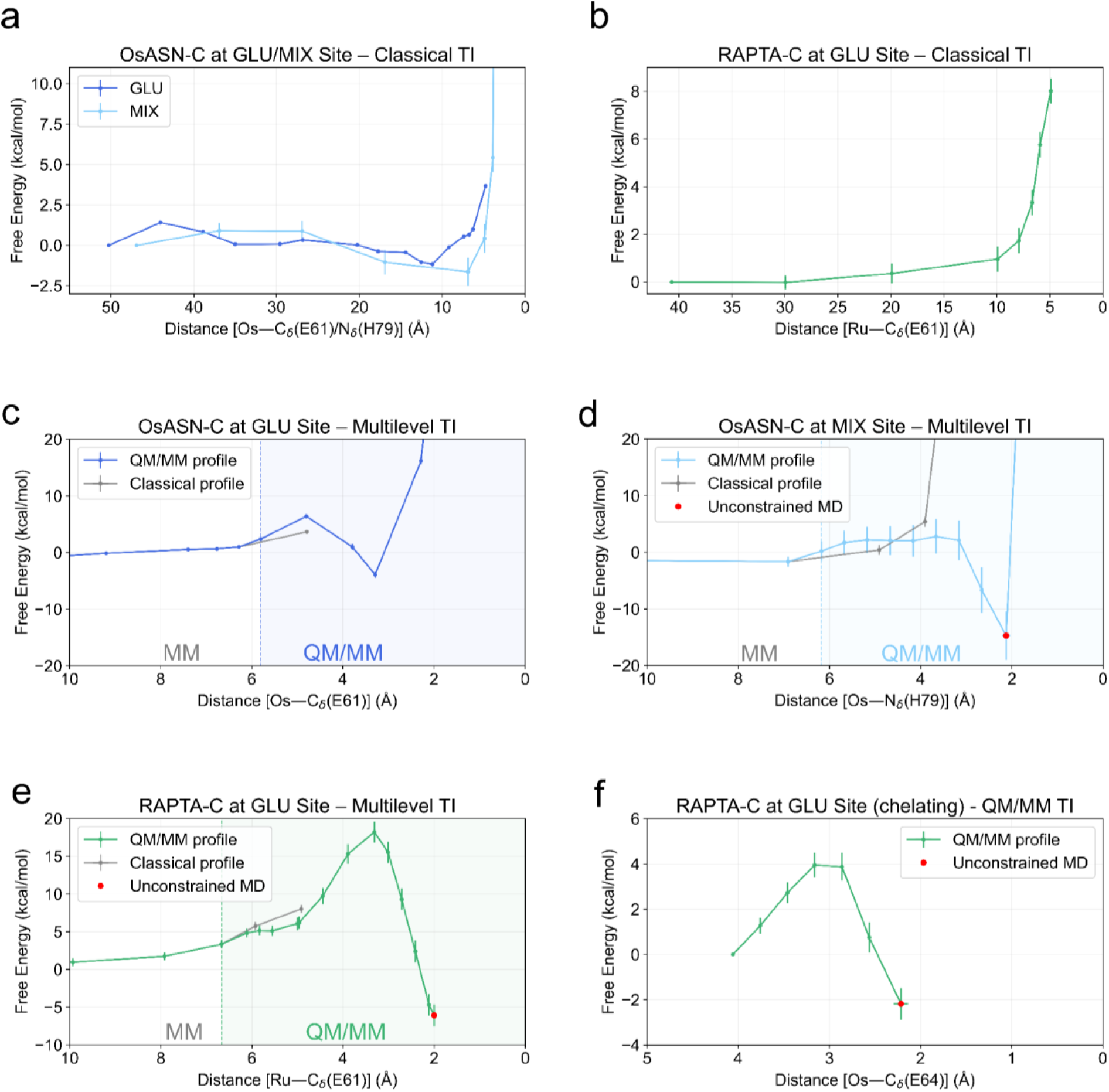
Free energy profiles for OsASN-C and RAPTA-C binding at the GLU and MIX sites. **a)** OsASN-C binding at the GLU and MIX sites, classical TI. **b)** RAPTA-C binding at the GLU site, classical TI. **c)** Multilevel free energy profile of binding at the GLU site for OsASN-C. **d)** Multilevel free energy profile of binding at the MIX site for OsASN-C. **e)** Multilevel free energy profile of binding at the GLU site for RAPTA-C. **f)** QM/MM free energy profile associated with the chelation at H2A E64 upon initial (monodentate) binding at H2A E61. The regions in c–e in which QM/MM TI is performed are highlighted with a shaded background.

From the MM-TI of the OsASN-C adduct at the GLU site (Os coordination at H2A E61), we could observe that the adduct shows a wide range of sampled conformations at this binding site. Hence, we analyzed the contacts that would result if both GLU and MIX binding sites were occupied simultaneously (**Figure 6**). This analysis indicates that an adduct at the MIX site would preclude the simultaneous presence of another adduct at the GLU site: a close contact analysis (more details provided in the Materials and Methods section) shows a large and widely oscillating number of 8.3±9.2 short atomic contacts ≤2.0 Å, with an average close contact distance of 1.3±0.8 Å. In contrast, the same analysis for QM/MM simulations of RAPTA-C reveals that a single adduct at the GLU site (Ru coordination at H2A E61 and H2A E64) is compatible with the simultaneous occupation of the MIX site: the close contact analysis does not show any close atomic contacts ≤2.0 Å, with an average close contact distance of 3.8±0.3 Å. This is in agreement with the experimental crystal structure, where both GLU and MIX sites are occupied for RAPTA, while for OsASN-C, only the MIX site is occupied.

**Figure 6.**
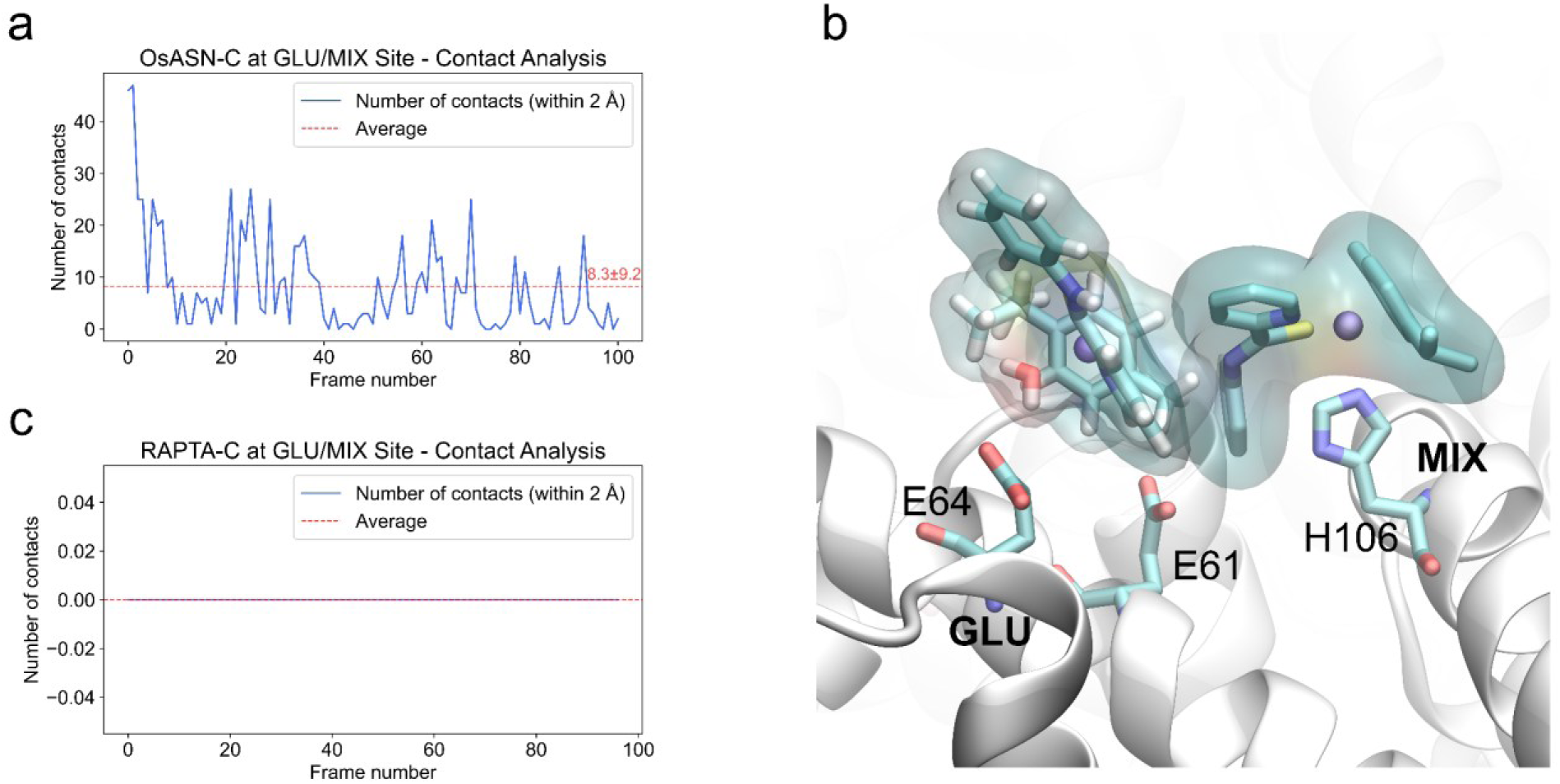
Analysis of inter-compound contacts when OsASN-C or RAPTA-C are occupying both the GLU and MIX sites simultaneously. The conformations of a compound at the GLU site were sampled with classical TI and analyzed with respect to the crystal structure having an OsASN-C adduct (**a**,**b**) or a RAPTA-C adduct (**c**) at the MIX site. **a)** OsASN-C close contact profile. The average number of contacts (8.3±9.2 short atomic contacts ≤2.0 Å) is shown in red. The average close contact distance is 1.3±0.8 Å. **b)** Representative snapshot from the classical TI at the minimum aligned to the crystal structure with the MIX site occupied, representing the van der Waals surfaces for the two compounds at the GLU and MIX sites. Hydrogen atoms are not present in the crystal structure but are represented for the compound at the GLU site (from the MD simulation). **c)** RAPTA-C contacts within 2.0 Å (none). The average close contact distance is 3.8±0.3 Å.

The facile and highly favorable energetics associated with OsASN-C binding at the MIX site are consistent with rapid coordination resulting in a highly stable adduct, which consequently sterically blocks binding of a second OsASN-C at the GLU site. On the other hand, RAPTA-C forms adducts at both the GLU and MIX sites that are mutually compatible. QM/MM-TI calculations illustrate the reaction profile of RAPTA-C binding to the GLU site, which is associated with an initial energetic hurdle for water ligand departure and coordination at H2A E61 (**Figure 5e**). However, subsequent chelation at H2A E64 is almost spontaneous (additional barrier of ∼4 kcal/mol, **Figure 5f**) and results in a substantial overall binding free energy of about −7 kcal/mol, which is in excellent agreement with our earlier mass spectrometry-based measurements of RAPTA-C binding to the NCP that yielded an overall association constant of 2.3×10^5^ M^−1^ (corresponding to a binding free energy of −7.3 kcal/mol; 28,29). Indeed, the bidentate character of the end-product apparently limits the conformational freedom of the GLU-RAPTA-C adduct, yielding an effective steric volume that does not preclude the binding of another RAPTA-C molecule to the adjacent MIX adduct (also confirmed by the contact analysis, **Figure 6**).

### Non-Leaving Ligand & Metal Substitutions Refine Site Selectivity

While the coordination and non-leaving ligand character of OsASN-C precludes concurrent binding at both the GLU and MIX sites, the favored adduct at the latter location coincides with the phenyl group of the SN ligand settling into a surface niche involving van der Waals and hydrophobic contacts with histone elements of H2A and H2B (**Figure 7a & 7b**). This added contact area contributes to the high stability of the final adduct (**Figure 5d**). Indeed, substitution of the metal center renders RuASN-C susceptible to loss of its SN ligand, and it is not observed to bind to the MIX site in the crystallographic analyses.

**Figure 7.**
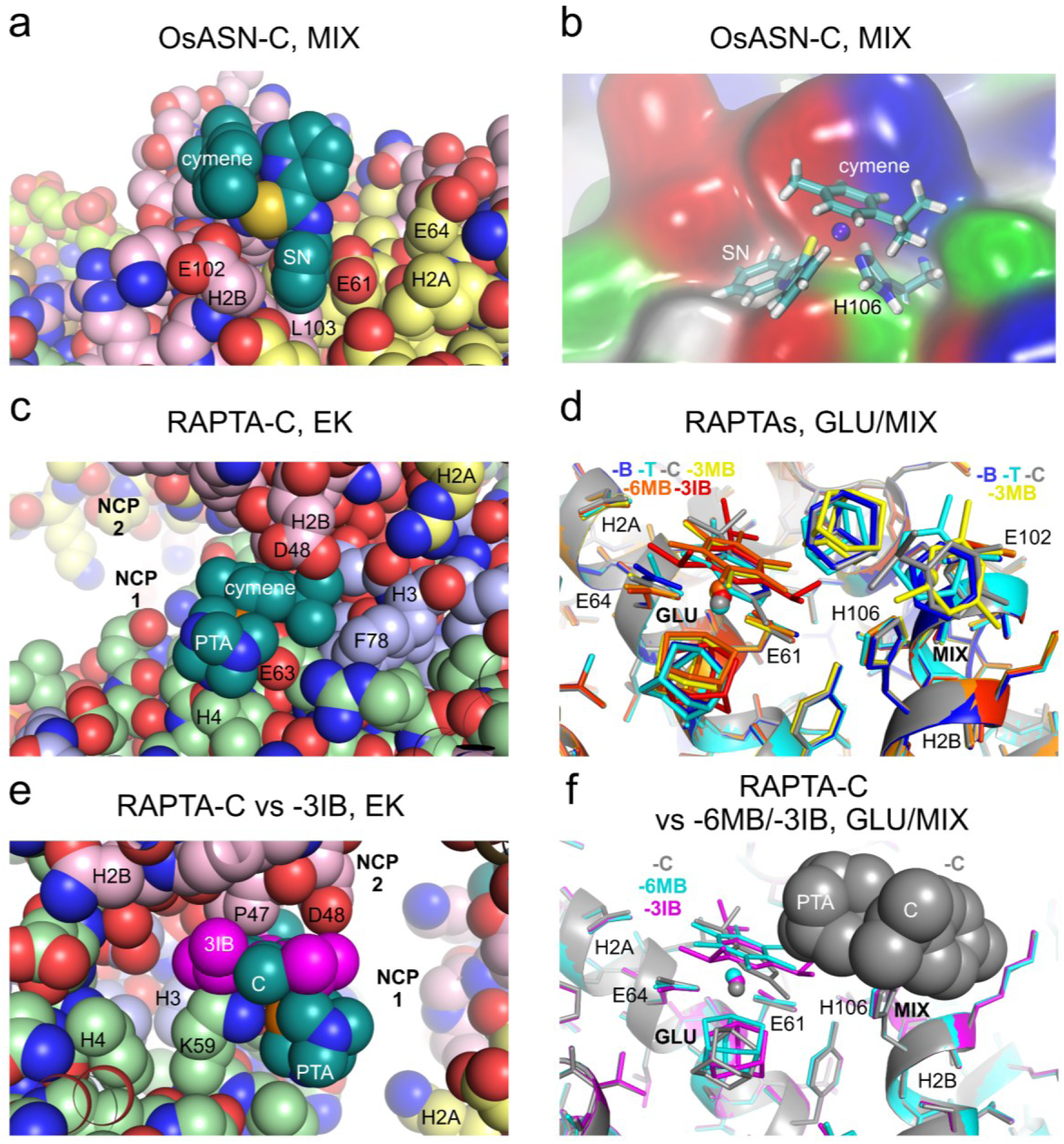
Non-leaving ligand shape complementarity and compatibility modulate histone site occupancy. **a,b)** Space-filling (a) and surface character (b) representations of OsASN-C binding at the MIX site (PDB ID 4J8U). The histone surface in (b) is coloured according to polarity: non-polar residues (white), basic residues (blue), acidic residues (red), and polar residues (green). **c,e)** Space-filling representations of RAPTA-C binding at the EK site, which is formed by a nucleosome–nucleosome (NCP1–NCP2) interface in the crystal. In (e), 1,3,5-triisopropyl-benzene (3IB) (magenta) has been superimposed onto the RAPTA-C cymene. **d)** Crystal structures showing RAPTA-B (blue), -T (cyan), -C (grey), -3MB (yellow), -6MB (orange), and -3IB (red) binding at the GLU and (for RAPTA-B/T/C/3MB) MIX sites. The six independent NCP structures are superimposed. **f)** The RAPTA-C (grey), RAPTA-6MB (cyan), and RAPTA-3IB (magenta) crystal structures illustrating the potential steric conflict between the arene of a GLU site-bound adduct with the PTA ligand of a MIX site-bound adduct (RAPTA-C shown in space-filling) when the arene is fully-substituted (6MB) or has multiple bulky substituents (3IB). The three independent NCP structures are superimposed.

While substitutions of the metal center can introduce or eliminate a particular histone binding site within the activity profile, we found that this can also be achieved through subtle alterations in the non-leaving ligand. In the first study of a RAPTA compound binding to chromatin, RAPTA-C was observed to bind to a third electronegative surface site on the nucleosome, the EK site, in addition to the GLU and MIX locations (28; **Figure 2a & 2e**). At this site, the Ru ion is coordinated to both K59 and E63 of H4, with the cymene engaged in further van der Waals contacts with adjacent H3 residues as well as those of H2B stemming from the neighboring nucleosome core particle (NCP) in the crystal (**Figure 7c**).

Interested in whether RAPTA agents with different arenes would display the same site selectivity, we investigated the nucleosome binding of a RAPTA family, in which the methyl and/or isopropyl groups of the cymene [C] are absent (benzene [B] and toluene [T]), or where the cymene is replaced by 1,3,5-trimethylbenzene [3MB], 1,3,5-triisopropylbenzene [3IB], or hexamethylbenzene [6MB]. Curiously, while they are all seen to bind to the GLU site, and at least to a certain extent also to the MIX site, none of the five non-cymene agents associates with the EK site in the NCP crystal structures (**Figure 7d** & **Tables 1, 2, and 4**). Structural analysis reveals that the RAPTA-C cymene isopropyl group is nestled within a significantly hydrophobic niche formed by the two stacking NCPs in the crystal (**Figure 7c**). This particular group is absent in all but one of the agents, RAPTA-3I, where the presence of the additional isopropyl groups at the ortho positions, however, likely presents a steric conflict with histone elements at the EK site (**Figure 7e**). While the increased substitutions on the arene ring, relative to cymene, associated with 3M and 6M may also pose steric challenges to binding at the EK site, this could not be the case for the B and T derivatives. Indeed, this reflects the importance of shape/hydrophobicity complementarity in adduct stability (site occupancy observed in the crystallographic analyses), as is seen for the Ru/OsASN-C compounds. RAPTA-B/T are lacking the additional isopropyl-histone contacts that are available to RAPTA-C, which, moreover, is not observed to bind at the symmetry-related EK’ site that is remote from any crystal contacts.

Regarding the simultaneous occupation of the GLU and MIX sites, the RAPTA-B, -T, -C, and -3M derivatives all display high occupancy binding at both locations in the crystal structures (**Figure 7d** and **Table 4**). However, for RAPTA-6M and RAPTA-3I, while there is a weak anomalous difference electron density signal at the MIX binding site indicative of low Ru occupancy, binding at the GLU site is the major adduct, which in turn sterically precludes or substantially diminishes binding at the adjacent MIX site (**Figure 7f**). Collectively, the identity of the arene ligand is seen to facilitate or diminish RAPTA binding in a highly arene-and histone-site-sensitive manner.

**Table 4.** Ruthenium(II) binding sites in NCP treated with the RAPTA agents.

| Agent | Anomalous Difference Peak Height ($\sigma$) | | |
| --- | --- | --- | --- |
|  | GLU | MIX | EK |
| RAPTA-B | 7.7 | 7.0 | -- |
| RAPTA-T | 6.8 | 6.1 | -- |
| RAPTA-C | 9.4 | 6.6 | 8.3 |
| RAPTA-3MB | 7.2 | 6.2 | -- |
| RAPTA-6MB | 5.6 | 3.4 | -- |
| RAPTA-3IB | 7.8 | 3.4 | -- |

Values correspond to anomalous difference electron density map peak magnitudes $>3.0\sigma$ at the GLU, MIX, and EK sites.

## DISCUSSION

With histone dysregulation materializing as a signature shared by both cancerous and aged cells, the histone proteins are emerging as multifunctional therapeutic targets (87-89). In a fashion analogous to how epigenetic drugs undermine cancer cell dependencies on specific chromatin-associated proteins, selective histone targeting could exploit cancer-specific vulnerabilities arising from disproportionate reliance on activity linked to a defined histone site, isoform, or variant. Our past work on Ru and Au complexes indicated a therapeutic (anticancer) potential in targeting selective histone sites in chromatin (28-30,32,33,90). Nevertheless, while metal-based agents can have certain advantages over purely organic small molecules, such as binding affinity and thus potency potential, achieving target selectivity is a leading challenge, and most metalloagents appear to associate with many different sites in vivo.

In an effort to develop more site-selective chromatin-targeting organometallic compounds, our objective here was to understand the basis for the differential histone binding preferences within and between the RAPTA and Ru/OsASN agents, which notably display distinct profiles in associating with nucleosomal surface and crevice sites. This would appear to be a crucial issue in determining the genomic activity of histone-targeting compounds, since binding to surface regions, such as the acidic patch, can modulate interactions with chromatin factors and neighboring nucleosomes, while binding at internal sites is more likely to influence nucleosome dynamics (assembly/disassembly) alone. We earlier found that, depending on the type of agent and therefore the binding site(s) occupied, histone adducts on the nucleosome surface can either facilitate, diminish, or block association of a chromatin-binding protein in vitro (32,33).

The strict histidine-coordinating activity of the Ru/OsASN compounds compared to the glutamate site preferences of the RAPTA agents implied initially that the nucleosomal surface (i.e., GLU, MIX, EK sites) versus crevice (i.e., HIS sites) discrimination was stemming from chemical/electronic differences. However, we find instead that the rigidity and spherical steric profile afforded by the arene and PTA non-leaving ligands in the RAPTA agents prevent this family of compounds from binding at the crevice sites (HIS/HIS’). On the other hand, the Ru/OsASN agents have a similar overall steric volume but with a flatter and more dynamic conformational character that allows them to coordinate at these internal locations. This suggests that even bulkier/larger non-leaving ligands could be employed to increase selectivity for the nucleosome crevice region (involving a greater histone contact area) by upholding a scaffold that can maintain or readily adopt a flat steric profile. In parallel, increasing ligand dimension can modulate binding activities at other locations.

Steric exclusion is also observed to dictate site occupancy even at the otherwise freely accessible surface locations when two sites are sufficiently close to compete with one another. This is the case for OsASN-C, where there is a favorable binding pocket associated with the MIX site that is nevertheless incompatible with simultaneous occupation at the adjacent GLU site. While OsASN-C adducts can apparently form at both sites, the much higher stability MIX adduct would dominate at equilibrium, rationalizing why only this site is seen to be occupied in the crystal structures. On the other hand, the bifunctional character and blunt steric profile of the RAPTA (i.e., -B/-T/-C/-3MB) compounds facilitate their concurrent coordination at both the GLU and MIX sites by fostering the formation of favorable inter-adduct van der Waals contacts between the arene and PTA ligands. However, for the RAPTA-6MB and -3IB agents, the high level of substitution and/or bulk of the arenes result in a steric conflict that favors occupation of only the GLU site.

Whereas steric exclusion factors associated with the identity of the non-leaving ligands are seen to operate in preventing otherwise potential binding at glutamate or histidine residues, the reverse— by virtue of favorable hydrophobic or van der Waals interactions— can be seen to yield additional coordination sites. This is observed at the levels of both metal and non-leaving ligand specifics. Notably, the substitution of Ru in place of Os substantially increases non-leaving ligand lability, which results in disfavoring RuASN-C binding at the MIX site. On the other hand, the identity of the arene can allow for a lock-and-key type of association, as seen for RAPTA coordination at the EK site. Here, both shape/hydrophobic complementarity as well as steric exclusion factors appear to be at play, depending on the exact nature of the arene ligand. The unique proclivity of RAPTA-C for the EK site, which is formed by a nucleosome-nucleosome stacking interaction within the NCP crystal, highlights the importance of the nucleosomal context in vivo, which can vary by genomic location and activity to block existing binding sites or create new ones (28).

In summary, a picture for histone adduct formation potential and stability emerges in which steric factors associated with interacting nucleosomes (i.e., chromatin) and individual nucleosomes serve as initial site discrimination filters (**Figure 8**). Metal identity and non-leaving ligand attributes modulate absolute and relative adduct stability, ultimately determining the resulting binding profiles. These principles could help in interpreting distinctions in therapeutic activities between closely related organometallic agents and for fine-tuning such compounds. For example, the dual nucleosome internal crevice/acidic patch-binder, OsASN-C, displays a distinct cancer cell line activity profile, consistent with disruption of DNA synthesis, that is not observed for the crevice-binding RuASN-C analogue (41). In a more general sense, the factors found to determine selectivity for the nucleosome should be applicable to protein-binding metal complexes in their preferential association to certain amino acid sites over others.

**Figure 8.**
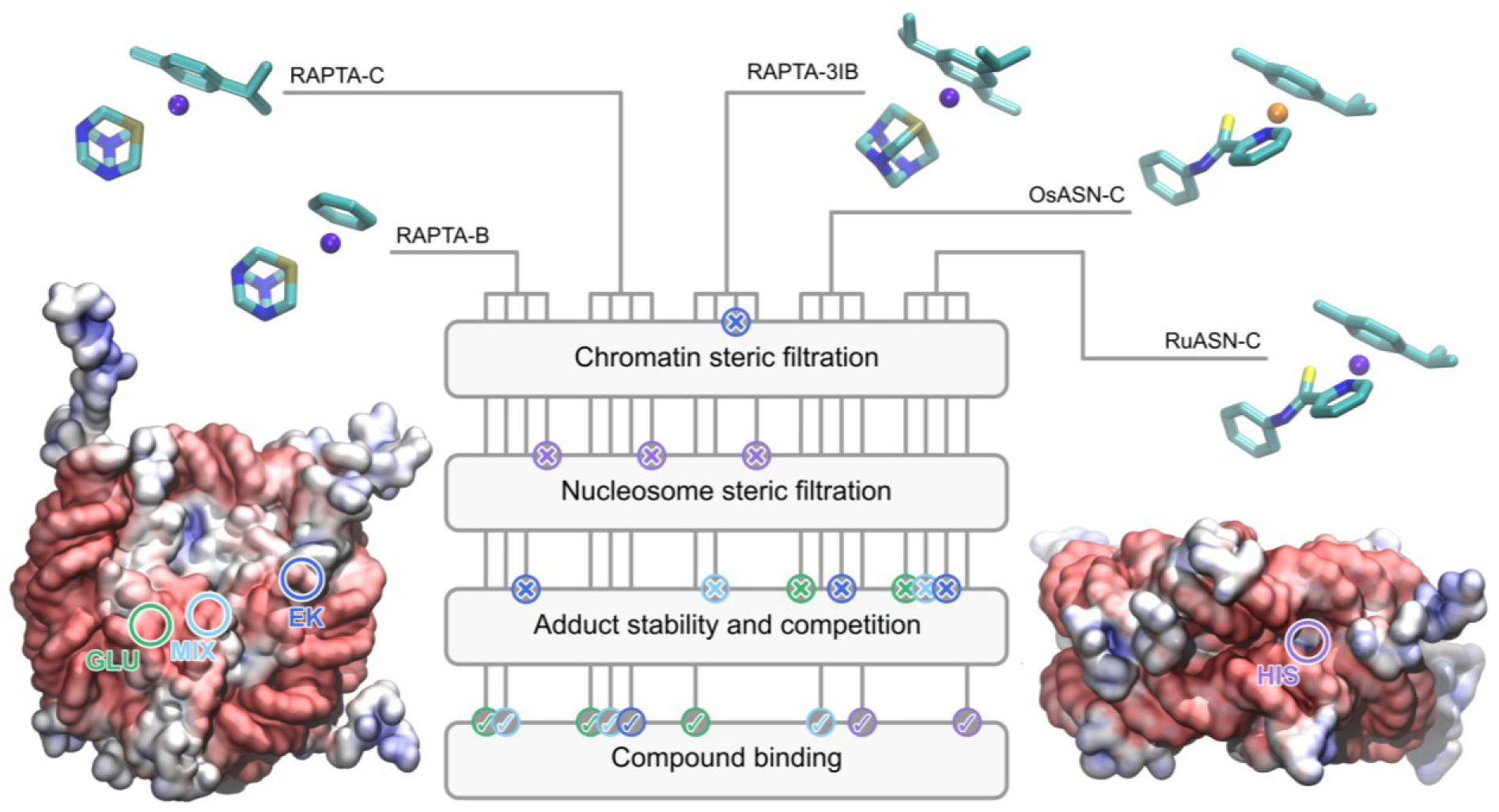
Proposed scheme of factors determining histone site selectivity in chromatin.

In our efforts to understand the basis of the distinct and overlapping nucleosome site selectivity profiles for Ru/Os arene compounds, we have developed a robust computational platform for the in-depth characterization of organometallic small molecule binding to nucleosomal systems. This involves combining experimentally-derived (crystallographic) site discrimination profiles and structures from compound screening with multilevel (MM, QM, QM/MM) simulations that are tailored to address the highly complex structural and energetic attributes. We believe this methodology promises to facilitate the characterization of compounds of interest and foster in silico-driven development of chromatin-targeting tools and therapeutic agents. In particular, our multilevel approach enables the deconvolution of empirically determined protein site selectivity features, guiding rational modifications to the metal center and non-leaving ligands in organometallic complexes.

## ACKNOWLEDGEMENTS

We thank V. Olieric, M. Wang, and staff for help at the Swiss Light Source (Paul Scherrer Institute, Villigen, Switzerland), M.S. Ong for assistance with X-ray data processing, and R. Muhammad for contributing to the X-ray data collection. Computational resources from the Swiss National Computing Centre CSCS are gratefully acknowledged.

## AUTHOR CONTRIBUTIONS

Computational experiments, A.L., T.E., G.P., U.R.; X-ray crystallography, Z.A., C.A.D.; Compound synthesis, A.A.N., C.G.H., P.J.D.; Data curation, analysis, validation, and visualization, A.L., U.R., C.A.D.; Conceptualization, funding acquisition, project administration, and supervision, P.J.D., U.R., C.A.D.; Writing‒ original draft, A.L., U.R., C.A.D.; Writing‒ review and editing, all authors.

## SUPPLEMENTARY DATA

Supplementary data are available at NAR online.

## CONFLICT OF INTEREST

None declared.

## FUNDING

This project was funded by the Polish National Science Center grant DEC-2024/53/B/NZ7/03477, the Singapore Ministry of Education Academic Research Fund Tier 2/3 (MOE-T2EP30121-0005, MOE2012-T3-1-001), and the Swiss National Science Foundation Grant Nos. 200020-185092 and 200020-219440.

## DATA AVAILABILITY

Atomic coordinates and structure factors for the crystal structures of NCP with bound RAPTA-B, -T, -C, -3MB, -6MB, and -3IB are deposited in the Protein Data Bank under accession codes 9RY3, 9RXV, 9RXU, 9RYA, 9RZK, and 9RZN, respectively. Data supporting the findings of this work are available within the article and Supplementary Information files. The computational data are available on Zenodo under https://zenodo.org/records/17258752, including Jupyter Notebooks and VMD visualization states to reproduce the analysis and generate the plots and figures presented.

## SUPPLEMENTARY DATA

**Table S1.**
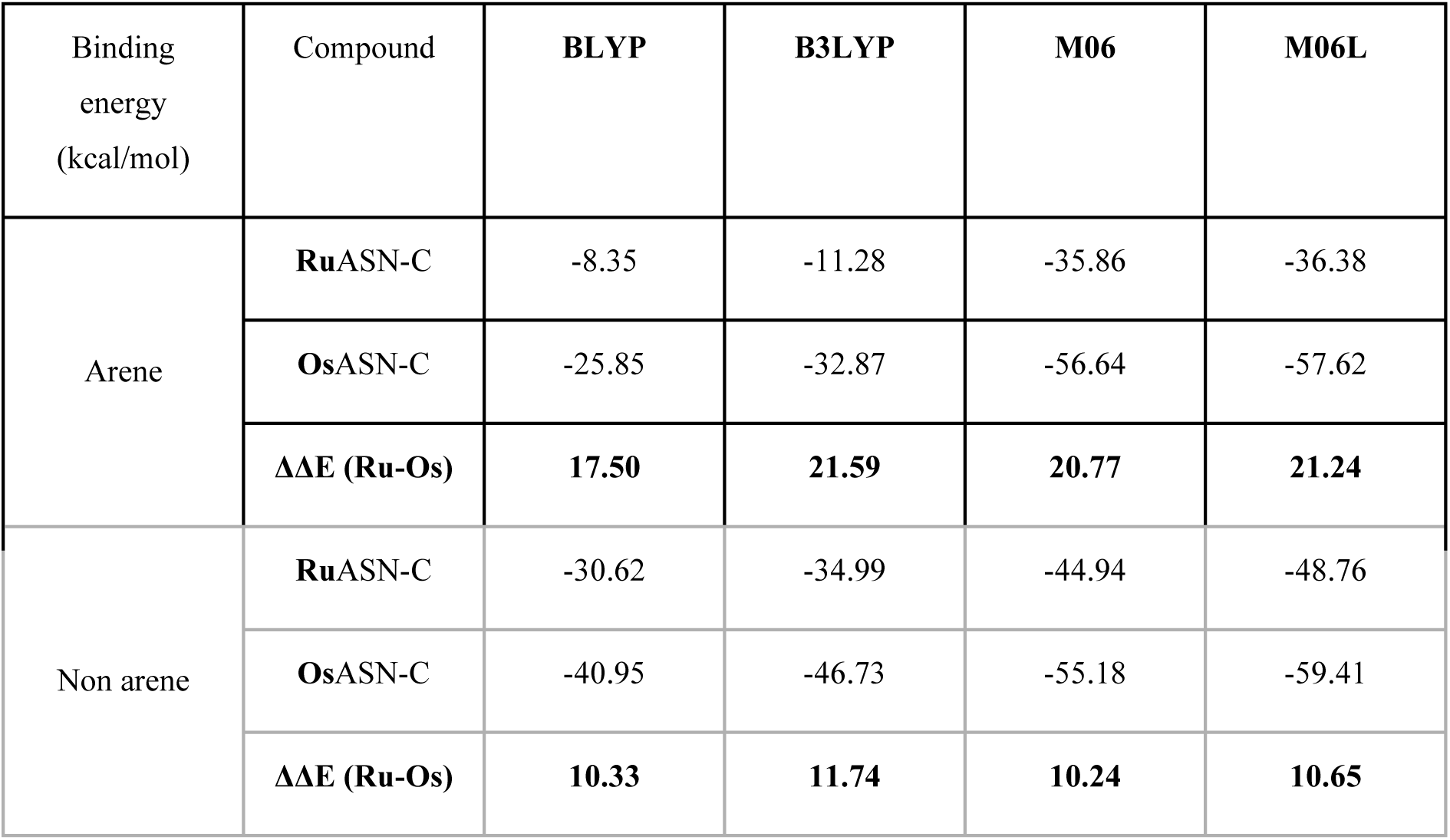
Binding energies for the arene and non-arene ligands in RuASN-C vs OsASN-C compounds obtained with different DFT functionals.

| Binding energy<br>(kcal/mol) | Compound | **BLYP** | **B3LYP** | **M06** | **M06L** |
| --- | --- | --- | --- | --- | --- |
| Arene | **RuASN-C** | -8.35 | -11.28 | -35.86 | -36.38 |
|  | **OsASN-C** | -25.85 | -32.87 | -56.64 | -57.62 |
|  | **$\Delta\Delta E$ (Ru-Os)** | **17.50** | **21.59** | **20.77** | **21.24** |
| Non arene | **RuASN-C** | -30.62 | -34.99 | -44.94 | -48.76 |
|  | **OsASN-C** | -40.95 | -46.73 | -55.18 | -59.41 |
|  | **$\Delta\Delta E$ (Ru-Os)** | **10.33** | **11.74** | **10.24** | **10.65** |

**Figure S1.**
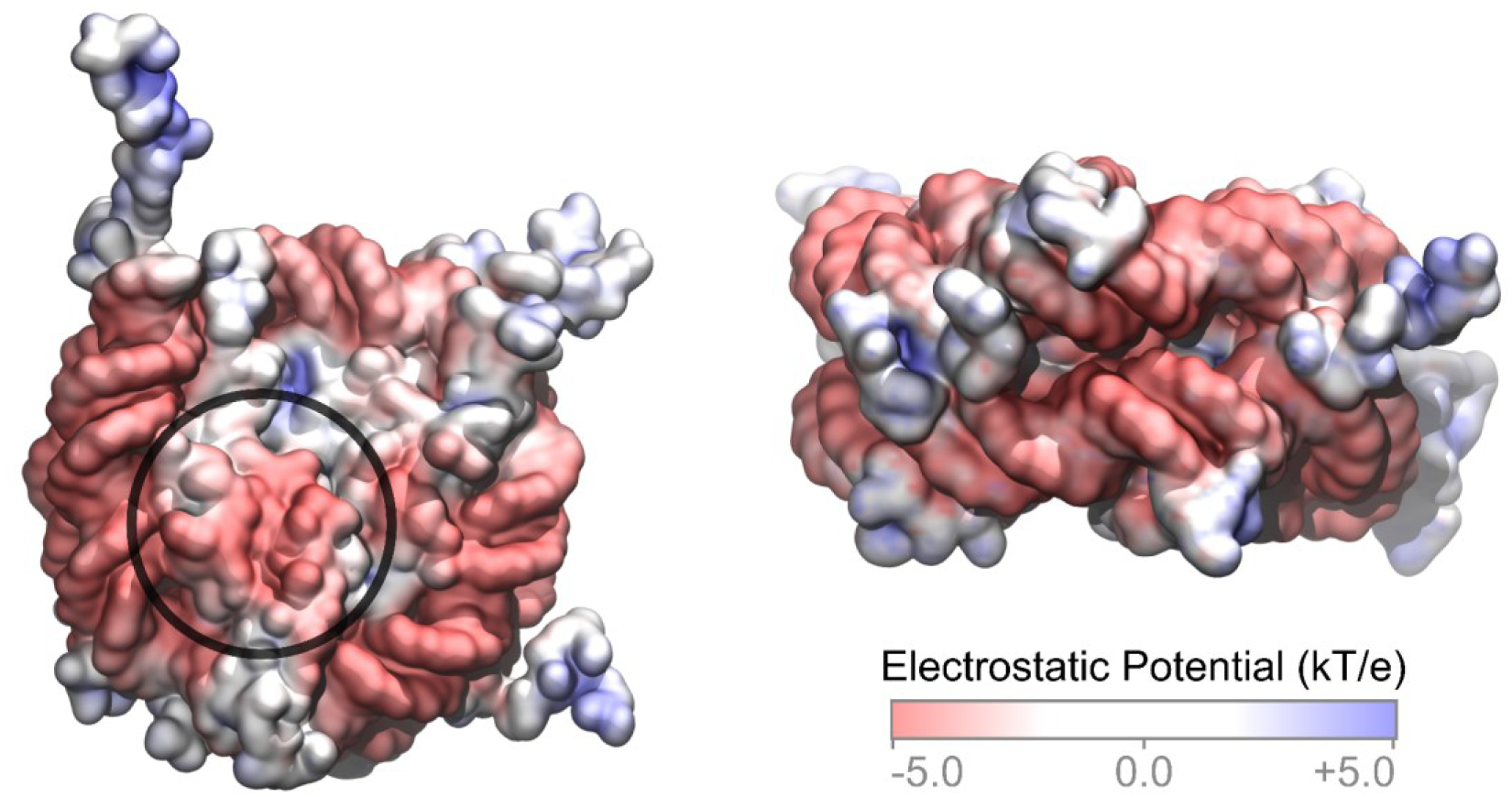
Electrostatic potential surface for the nucleosome core particle (NCP; PDB ID 1KX5; 39) computed with APBS VMD plugin, after protonation with H++ server (http://newbiophysics.cs.vt.edu/H++/). The acidic patch is highlighted with a black circle.

**Figure S2.**
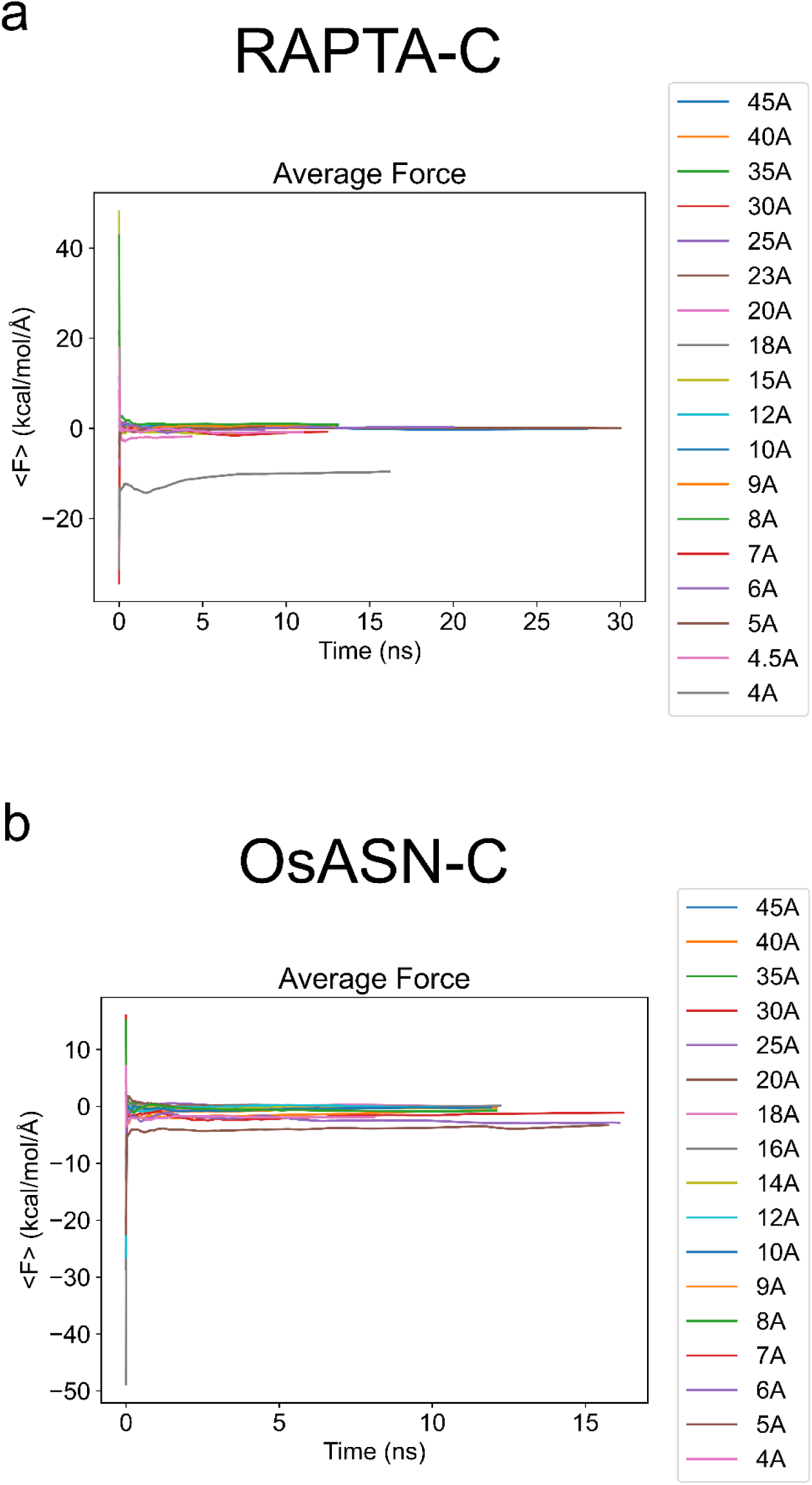
Convergence of the average force in the classical TI of the binding to the HIS site for RAPTA-C (**a**) and OsASN-C (**b**).

**Figure S3.**
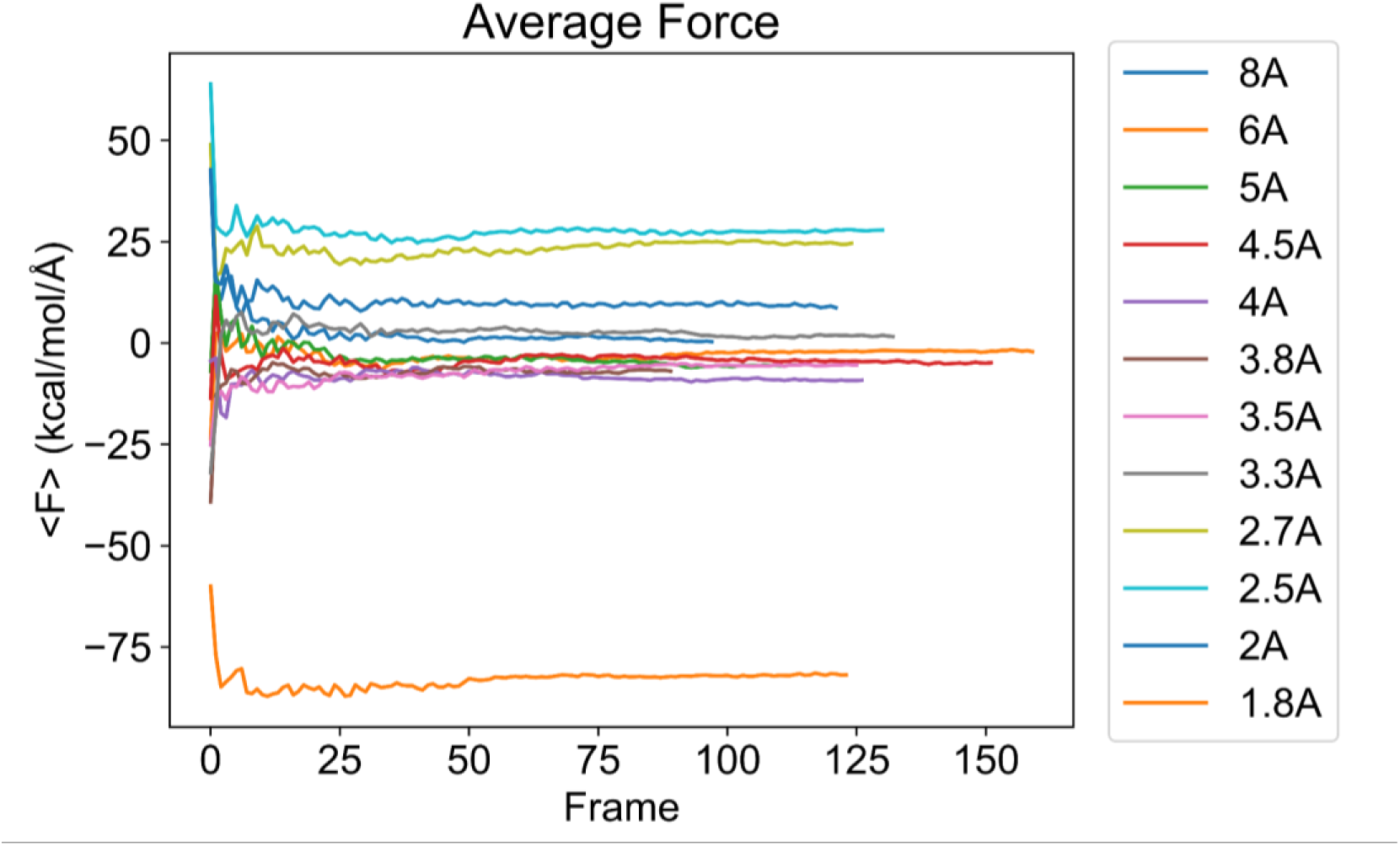
Convergence of the average force in the QM/MM TI of the binding to the HIS site for OsASN-C.

**Figure S4.**
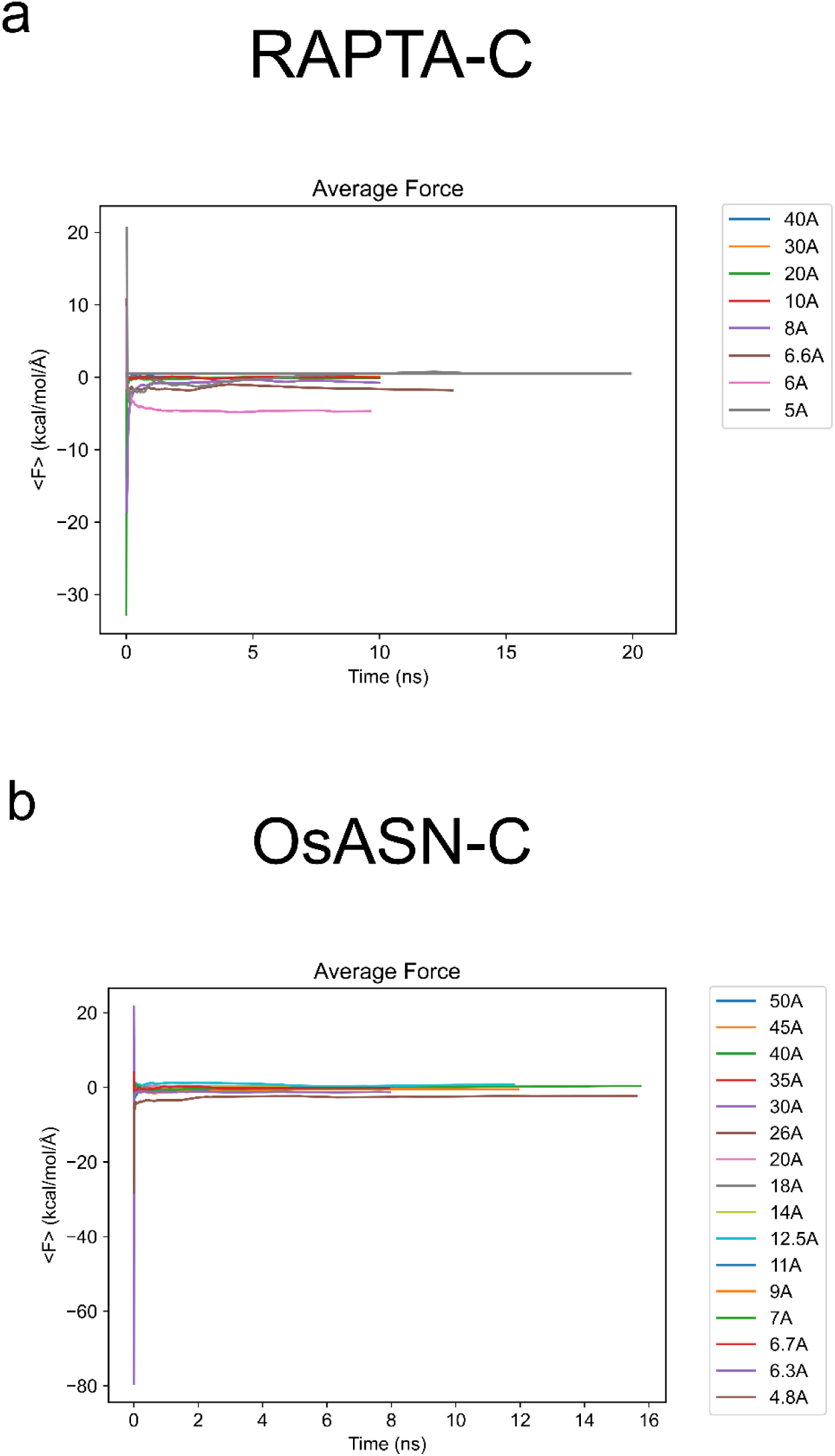
Convergence of the average force in the classical TI of the binding to the GLU site for RAPTA-C (a) and OsASN-C (b).

**Figure S5.**
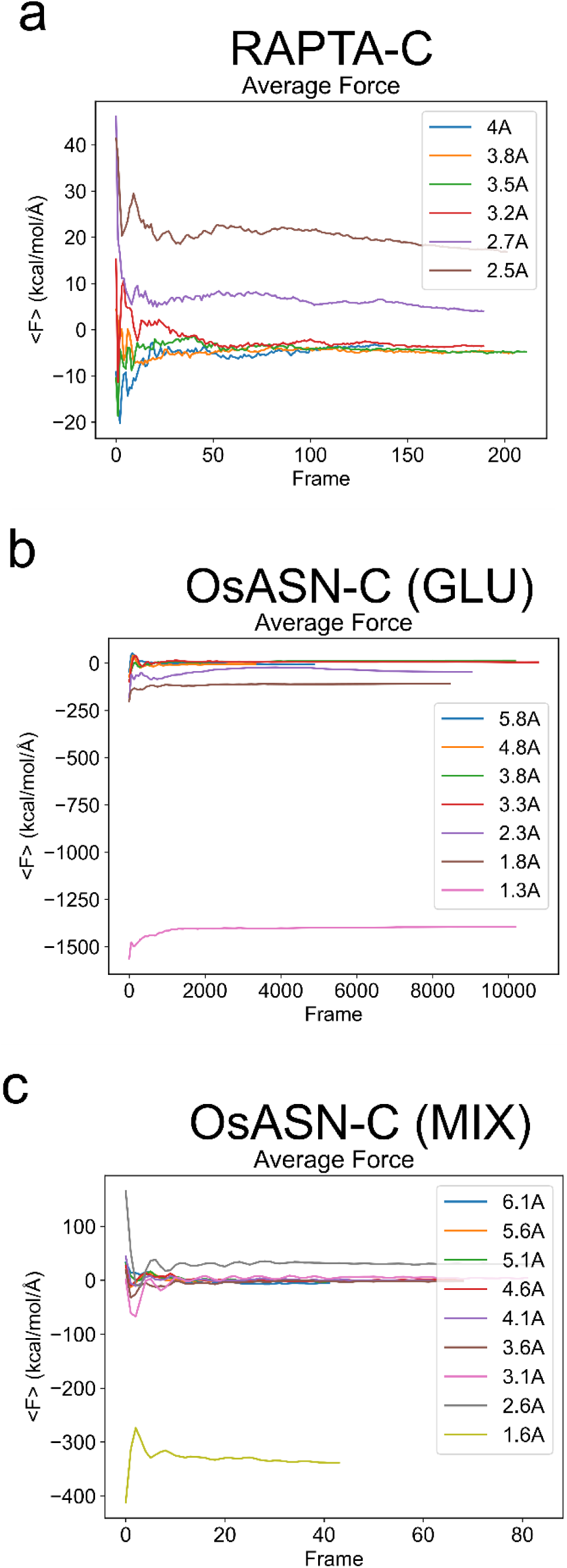
Convergence of the average force in the QM/MM TI of the binding for RAPTA-C to the GLU site (a), OsASN-C to the GLU site (b), and OsASN-C to the MIX site (c).

## Notes

https://doi.org/10.5281/zenodo.17258752

### Competing Interest Statement

The authors have declared no competing interest.

### Summary of Updates

The manuscript has been substantially reorganized. Several results previously in the Supplementary Information have been moved into the main text to improve clarity and accessibility.

## REFERENCES

1. Soldevila-Barreda, J.J. and Metzler-Nolte, N. Intracellular Catalysis with Selected Metal Complexes and Metallic Nanoparticles: Advances toward the Development of Catalytic Metallodrugs. Chem Rev 2019;119:829–869. 10.1021/acs.chemrev.8b00493

2. Vigueras, G. and Gasser, G. Anticancer platinum-based photo-oxidants in a new light. Nat Chem 2023;15:896–898. 10.1038/s41557-023-01250-w

3. Chang, M.R., Rusanov, D.A., Arakelyan, J., Alshehri, M., Asaturova, A., Kireeva, G.S., Babak, M. and Ang, W.H. Targeting emerging cancer hallmarks by transition metal complexes: Cancer stem cells and tumor microbiome. Part I. Coord Chem Rev 2023;477:214923. 10.1016/j.ccr.2022.214923

4. Rottenberg, S., Disler, C. and Perego, P. The rediscovery of platinum-based cancer therapy. Nat Rev Cancer 2021;21:37–50. 10.1038/s41568-020-00308-y

5. Alassadi, S., Pisani, M.J. and Wheate, N.J. A chemical perspective on the clinical use of platinum-based anticancer drugs. Dalton Trans 2022;51:10835–10846. 10.1039/d2dt01875f

6. Wu, B., Davey, G.E., Nazarov, A.A., Dyson, P.J. and Davey, C.A. Specific DNA structural attributes modulate platinum anticancer drug site selection and cross-link generation. Nucleic Acids Res 2011;39:8200–8212. 10.1093/nar/gkr491

7. Oun, R., Moussa, Y.E. and Wheate, N.J. The side effects of platinum-based chemotherapy drugs: a review for chemists. Dalton Trans 2018;47:6645–6653. 10.1039/c8dt00838h

8. Merlino, A. Interactions between proteins and Ru compounds of medicinal interest: A structural perspective. Coord Chem Rev 2016;326:111–134. 10.1016/j.ccr.2016.08.001

9. Allardyce, C.S. and Dyson, P.J. Metal-based drugs that break the rules. Dalton Transactions 2016;45:3201–3209. 10.1039/c5dt03919c

10. Alessio, E. and Messori, L. NAMI-A and KP1019/1339, Two Iconic Ruthenium Anticancer Drug Candidates Face-to-Face: A Case Story in Medicinal Inorganic Chemistry. Molecules 2019;24:1995. 10.3390/molecules24101995

11. Murray, B.S. and Dyson, P.J. Recent progress in the development of organometallics for the treatment of cancer. Curr Opin Chem Biol 2020;56:28–34. 10.1016/j.cbpa.2019.11.001

12. Happl, B., Balber, T., Heffeter, P., Denk, C., Welch, J.M., Koester, U., Alliot, C., Bonraisin, A.C., Brandt, M., Haddad, F. et al. Synthesis and preclinical evaluation of BOLD-100 radiolabeled with ruthenium-97 and ruthenium-103. Dalton Transactions 2024;53:6031–6040. 10.1039/d4dt00118d

13. Murray, B.S., Babak, M.V., Hartinger, C.G. and Dyson, P.J. The development of RAPTA compounds for the treatment of tumors. Coord Chem Rev 2016;306:86–114. 10.1016/j.ccr.2015.06.014

14. Bolitho, E.M., Bridgewater, H.E., Needham, R.J., Coverdale, J.P.C., Quinn, P.D., Sanchez-Cano, C. and Sadler, P.J. Elemental mapping of half-sandwich azopyridine osmium arene complexes in cancer cells. Inorganic Chemistry Frontiers 2021;8:3675–3685. 10.1039/d1qi00512j

15. Xue, X.L., Fu, Y., He, L., Salassa, L., He, L.F., Hao, Y.Y., Koh, M.J., Soulié, C., Needham, R.J., Habtemariam, A. et al. Photoactivated Osmium Arene Anticancer Complexes. Inorg Chem 2021;60:17450–17461. 10.1021/acs.inorgchem.1c00241

16. Swaminathan, S., Deepak, R.J. and Karvembu, R. Interweaving catalysis and cancer using Ru- and Os-arene complexes to alter cellular redox state: A structure-activity relationship (SAR) review. Coord Chem Rev 2023;491:1–20. 10.1016/j.ccr.2023.215230

17. Yang, Q.Y., Ma, R., Gu, Y.Q., Xu, X.F., Chen, Z.F. and Liang, H. Arene-Ruthenium(II)/Osmium(II) Complexes Potentiate the Anticancer Efficacy of Metformin via Glucose Metabolism Reprogramming. Angewandte Chemie-International Edition 2022;61:6911–6919. 10.1002/anie.202208570

18. Riaz, Z., Lee, B.Y.T., Stjärnhage, J., Movassaghi, S., Söhnel, T., Jamieson, S.M.F., Shaheen, M.A., Hanif, M. and Hartinger, C.G. Anticancer Ru and Os complexes of N-(4-chlorophenyl) pyridine-2-carbothioamide: Substitution of the labile chlorido ligand with phosphines. J Inorg Biochem 2023;241:112115. 10.1016/j.jinorgbio.2022.112115

19. Pagliaricci, N., Pettinari, R., Marchetti, F., Pagliaricci, S., Cuccioloni, M., Eleuteri, A.M., Galindo, A., Fadaei-Tirani, F., Glinkina, K. and Dyson, P.J. Half-Sandwich Ruthenium and Osmium Complexes with Hydrazinocurcuminoid-like Ligands. Organometallics 2025;44:1155–1164. 10.1021/acs.organomet.5c00082

20. Baldi, S., Korber, P. and Becker, P.B. Beads on a string-nucleosome array arrangements and folding of the chromatin fiber. Nat Struct Mol Biol 2020;27:109–118. 10.1038/s41594-019-0368-x

21. McGinty, R.K. and Tan, S. Nucleosome structure and function. Chem Rev 2015;115:2255-2273. 10.1021/cr500373h

22. Nacev, B.A., Feng, L., Bagert, J.D., Lemiesz, A.E., Gao, J., Soshnev, A.A., Kundra, R., Schultz, N., Muir, T.W. and Allis, C.D. The expanding landscape of ‘oncohistone’ mutations in human cancers. Nature 2019;567:473–478. 10.1038/s41586-019-1038-1

23. Ferrand, J., Rondinelli, B. and Polo, S.E. Histone Variants: Guardians of Genome Integrity. Cells 2020;9:2424. 10.3390/cells9112424

24. Espiritu, D., Gribkova, A.K., Gupta, S., Shaytan, A.K. and Panchenko, A.R. Molecular Mechanisms of Oncogenesis through the Lens of Nucleosomes and Histones. J Phys Chem B 2021;125:3963–3976. 10.1021/acs.jpcb.1c00694

25. Bonner, E.R., Dawood, A., Gordish-Dressman, H., Eze, A., Bhattacharya, S., Yadavilli, S., Mueller, S., Waszak, S.M. and Nazarian, J. Pan-cancer atlas of somatic core and linker histone mutations. NPJ Genom Med 2023;8:23. 10.1038/s41525-023-00367-8

26. Wu, B., Droge, P. and Davey, C.A. Site selectivity of platinum anticancer therapeutics. Nat Chem Biol 2008;4:110–112. 10.1038/nchembio.2007.58

27. Chua, E.Y., Davey, G.E., Chin, C.F., Droge, P., Ang, W.H. and Davey, C.A. Stereochemical control of nucleosome targeting by platinum-intercalator antitumor agents. Nucleic Acids Res 2015;43:5284–5296. 10.1093/nar/gkv356

28. Wu, B., Ong, M.S., Groessl, M., Adhireksan, Z., Hartinger, C.G., Dyson, P.J. and Davey, C.A. A ruthenium antimetastasis agent forms specific histone protein adducts in the nucleosome core. Chemistry 2011;17:3562–3566. 10.1002/chem.201100298

29. Adhireksan, Z., Davey, G.E., Campomanes, P., Groessl, M., Clavel, C.M., Yu, H., Nazarov, A.A., Yeo, C.H., Ang, W.H., Droge, P. et al. Ligand substitutions between ruthenium-cymene compounds can control protein versus DNA targeting and anticancer activity. Nat Commun 2014;5:3462. 10.1038/ncomms4462

30. Davey, G.E., Adhireksan, Z., Ma, Z., Riedel, T., Sharma, D., Padavattan, S., Rhodes, D., Ludwig, A., Sandin, S., Murray, B.S. et al. Nucleosome acidic patch-targeting binuclear ruthenium compounds induce aberrant chromatin condensation. Nat Commun 2017;8:1575. 10.1038/s41467-017-01680-4

31. Ma, Z., Palermo, G., Adhireksan, Z., Murray, B.S., von Erlach, T., Dyson, P.J., Rothlisberger, U. and Davey, C.A. An Organometallic Compound which Exhibits a DNA Topology-Dependent One-Stranded Intercalation Mode. Angew Chem Int Ed Engl 2016;55:7441–7444. 10.1002/anie.201602145

32. Adhireksan, Z., Palermo, G., Riedel, T., Ma, Z., Muhammad, R., Rothlisberger, U., Dyson, P.J. and Davey, C.A. Allosteric cross-talk in chromatin can mediate drug-drug synergy. Nat Commun 2017;8:14860. 10.1038/ncomms14860

33. Batchelor, L.K., De Falco, L., von Erlach, T., Sharma, D., Adhireksan, Z., Roethlisberger, U., Davey, C.A. and Dyson, P.J. Crosslinking Allosteric Sites on the Nucleosome. Angew Chem Int Ed Engl 2019;58:15660–15664. 10.1002/anie.201906423

34. Meier, S.M., Hanif, M., Adhireksan, Z., Pichler, V., Novak, M., Jirkovsky, E., Jakupec, M.A., Arion, V.B., Davey, C.A., Keppler, B.K. et al. Novel metal(II) arene 2-pyridinecarbothioamides: a rationale to orally active organometallic anticancer agents. Chemical Science 2013;4:1837–1846. 10.1039/C3sc22294b

35. Scolaro, C., Bergamo, A., Brescacin, L., Delfino, R., Cocchietto, M., Laurenczy, G., Geldbach, T.J., Sava, G. and Dyson, P.J. In vitro and in vivo evaluation of ruthenium(II)-arene PTA complexes. J Med Chem 2005;48:4161–4171. 10.1021/jm050015d

36. Weiss, A., Berndsen, R.H., Dubois, M., Müller, C., Schibli, R., Griffioen, A.W., Dyson, P.J. and Nowak-Sliwinska, P. Anti-tumor activity of the organometallic ruthenium(II)-arene complex [Ru(η-cymene)-Cl(pta)] (RAPTA-C) in human ovarian and colorectal carcinomas. Chemical Science 2014;5:4742–4748. 10.1039/c4sc01255k

37. Nowak-Sliwinska, P., van Beijnum, J.R., Casini, A., Nazarov, A.A., Wagnieres, G., van den Bergh, H., Dyson, P.J. and Griffioen, A.W. Organometallic Ruthenium(II) Arene Compounds with Antiangiogenic Activity. J Med Chem 2011;54:3895–3902. 10.1021/jm2002074

38. Lee, R.F.S., Escrig, S., Croisier, M., Clerc-Rosset, S., Knott, G.W., Meibom, A., Davey, C.A., Johnsson, K. and Dyson, P.J. NanoSIMS analysis of an isotopically labelled organometallic ruthenium(II) drug to probe its distribution and state. Chem Commun 2015;51:16486–16489. 10.1039/c5cc06983a

39. Davey, C.A., Sargent, D.F., Luger, K., Maeder, A.W. and Richmond, T.J. Solvent mediated interactions in the structure of the nucleosome core particle at 1.9 Å resolution. J Mol Biol 2002;319:1097–1113. 10.1016/S0022-2836(02)00386-8

40. Meier, S.M., Kreutz, D., Winter, L., Klose, M.H.M., Cseh, K., Weiss, T., Bileck, A., Alte, B., Mader, J.C., Jana, S. et al. An Organoruthenium Anticancer Agent Shows Unexpected Target Selectivity For Plectin. Angewandte Chemie-International Edition 2017;56:8267-8271. 10.1002/anie.201702242

41. Klose, M.H.M., Schöberl, A., Heffeter, P., Berger, W., Hartinger, C.G., Koellensperger, G., Meier-Menches, S.M. and Keppler, B.K. Serum-binding properties of isosteric ruthenium and osmium anticancer agents elucidated by SEC-ICP-MS. Monatsh Chem 2018;149:1719–1726. 10.1007/s00706-018-2280-1

42. Lovett, J.H., Lai, B.P., Bloomfield, H.O., Baker, A.T., Sullivan, M.P., Hartinger, C.G. and Harris, H.H. X-ray fluorescence microscopy and X-ray absorption spectroscopy reveal the stability of the plecstatin-1 scaffold in biological model systems: comparison of Ru, Os and Ir analogues. Chemical Science 2025;16:11347–11358. 10.1039/d5sc02925b

43. Allardyce, C.S., Dyson, P.J., Ellis, D.J. and Heath, S.L. [Ru(eta(6)-p-cymene)Cl-2(pta)] (pta=1,3,5-triaza-7-phosphatricyclo[3.3.1.1]decane): a water soluble compound that exhibits pH dependent DNA binding providing selectivity for diseased cells. Chem Commun (Camb) 2001;15:1396-1397. 10.1039/B104021A

44. Ong, M.S., Richmond, T.J. and Davey, C.A. DNA stretching and extreme kinking in the nucleosome core. J Mol Biol 2007;368:1067–1074. 10.1016/j.jmb.2007.02.062

45. Leslie, A.G. The integration of macromolecular diffraction data. Acta Crystallogr D Biol Crystallogr 2006;62:48–57. 10.1107/S090744499900846X

46. Evans, P. Scaling and assessment of data quality. Acta Crystallogr D Biol Crystallogr 2006;62:72–82. 10.1107/S0907444905036693

47. Winn, M.D., Ballard, C.C., Cowtan, K.D., Dodson, E.J., Emsley, P., Evans, P.R., Keegan, R.M., Krissinel, E.B., Leslie, A.G., McCoy, A. et al. Overview of the CCP4 suite and current developments. Acta Crystallogr D Biol Crystallogr 2011;67:235–242. 10.1107/S0907444910045749

48. Agirre, J., Atanasova, M., Bagdonas, H., Ballard, C.B., Basle, A., Beilsten-Edmands, J., Borges, R.J., Brown, D.G., Burgos-Marmol, J.J., Berrisford, J.M. et al. The CCP4 suite: integrative software for macromolecular crystallography. Acta Crystallogr D Struct Biol 2023;79:449–461. 10.1107/S2059798323003595

49. Emsley, P., Lohkamp, B., Scott, W.G. and Cowtan, K. Features and development of Coot. Acta Crystallogr D Biol Crystallogr 2010;66:486-501. 10.1107/S0907444910007493

50. Murshudov, G.N., Vagin, A.A. and Dodson, E.J. Refinement of macromolecular structures by the maximum-likelihood method. Acta Crystallogr D Biol Crystallogr 1997;53:240–255. 10.1107/S0907444996012255

51. Vock, C.A., Renfrew, A.K., Scopelliti, R., Juillerat-Jeanneret, L. and Dyson, P.J. Influence of the diketonato ligand on the cytotoxicities of [Ru(η6-p-cymene)-(R2acac)(PTA)]+ complexes (PTA = 1,3,5-triaza-7-phosphaadamantane). Eur J Inorg Chem 2008;10:1661-1671. 10.1002/ejic.200701291

52. Levy, A., Slama, V., Guilbert, S., Antalík, A., Johnson, S.K., Frisari, G. and Rothlisberger, U. Modeling chemical reactivity in complex systems: Insights from hybrid QM/MM MD simulations. J Catal 2026;453. 10.1016/j.jcat.2025.116520

53. Luger, K., Mader, A.W., Richmond, R.K., Sargent, D.F. and Richmond, T.J. Crystal structure of the nucleosome core particle at 2.8 angstrom resolution. Nature 1997;389:251–260. 10.1038/38444

54. Maier, J.A., Martinez, C., Kasavajhala, K., Wickstrom, L., Hauser, K.E. and Simmerling, C. ff14SB: Improving the Accuracy of Protein Side Chain and Backbone Parameters from ff99SB. Journal of Chemical Theory and Computation 2015;11:3696–3713. 10.1021/acs.jctc.5b00255

55. Tian, C., Kasavajhala, K., Belfon, K.A.A., Raguette, L., Huang, H., Migues, A.N., Bickel, J., Wang, Y.Z., Pincay, J., Wu, Q. et al. ff19SB: Amino-Acid-Specific Protein Backbone Parameters Trained against Quantum Mechanics Energy Surfaces in Solution. Journal of Chemical Theory and Computation 2020;16:528–552. 10.1021/acs.jctc.9b00591

56. Perez, A., Marchan, I., Svozil, D., Sponer, J., Cheatham, T.E., 3rd, Laughton, C.A. and Orozco, M. Refinement of the AMBER force field for nucleic acids: improving the description of alpha/gamma conformers. Biophys J 2007;92:3817–3829. 10.1529/biophysj.106.097782

57. Jorgensen, W.L., Chandrasekhar, J., Madura, J.D., Impey, R.W. and Klein, M.L. Comparison of Simple Potential Functions for Simulating Liquid Water. J Chem Phys 1983;79:926–935. Doi 10.1063/1.445869

58. Gossens, C., Tavernelli, I. and Rothlisberger, U. DNA structural distortions induced by ruthenium-arene anticancer compounds. Journal of the American Chemical Society 2008;130:10921–10928. 10.1021/ja800194a

59. Desoize, B. Metals and metal compounds in cancer treatment. Anticancer Res 2004;24:1529–1544.

60. Scolaro, C., Hartinger, C.G., Allardyce, C.S., Keppler, B.K. and Dyson, P.J. Hydrolysis study of the bifunctional antitumour compound RAPTA-C, [Ru(η-cymene)Cl(pta). J Inorg Biochem 2008;102:1743–1748. 10.1016/j.jinorgbio.2008.05.004

61. Abraham, M.J., Murtola, T., Schulz, R., Páll, S., Smith, J.C., Hess, B. and Lindahl, E. GROMACS: High performance molecular simulations through multi-level parallelism from laptops to supercomputers. SoftwareX 2015;1:19–25. 10.1016/j.softx.2015.06.001

62. Van der Spoel, D., Lindahl, E., Hess, B., Groenhof, G., Mark, A.E. and Berendsen, H.J.C. GROMACS: Fast, flexible, and free. J Comput Chem 2005;26:1701-1718. 10.1002/jcc.20291

63. Berendsen, H.J.C., Vanderspoel, D. and Vandrunen, R. Gromacs - a Message-Passing Parallel Molecular-Dynamics Implementation. Comput Phys Commun 1995;91:43–56. 10.1016/0010-4655(95)00042-E

64. CPMD, Copyright IBM Corp 1990–2008; Copyright MPI für Festkörperforschung Stuttgart 1997–2001. Available at: https://doi.org/github.com/CPMD-code

65. Laio, A., VandeVondele, J. and Rothlisberger, U. A Hamiltonian electrostatic coupling scheme for hybrid Car-Parrinello molecular dynamics simulations. J Chem Phys 2002;116:6941–6947. 10.1063/1.1462041

66. Becke, A.D. Density-Functional Exchange-Energy Approximation with Correct Asymptotic-Behavior. Physical Review A 1988;38:3098–3100. 10.1103/PhysRevA.38.3098

67. Lee, C.T., Yang, W.T. and Parr, R.G. Development of the Colle-Salvetti Correlation-Energy Formula into a Functional of the Electron-Density. Physical Review B 1988;37:785–789. 10.1103/PhysRevB.37.785

68. Troullier, N. and Martins, J.L. Efficient Pseudopotentials for Plane-Wave Calculations. Physical Review B 1991;43:1993–2006. 10.1103/PhysRevB.43.1993

69. von Lilienfeld, O.A., Tavernelli, I., Rothlisberger, U. and Sebastiani, D. Optimization of effective atom centered potentials for London dispersion forces in density functional theory - art. no. 153004. Phys Rev Lett 2004;93. 10.1103/PhysRevLett.93.153004

70. Nose, S. A Unified Formulation of the Constant Temperature Molecular-Dynamics Methods. J Chem Phys 1984;81:511–519. 10.1063/1.447334

71. Hoover, W.G. Canonical Dynamics - Equilibrium Phase-Space Distributions. Physical Review A 1985;31:1695–1697. 10.1103/PhysRevA.31.1695

72. Carter, E.A., Ciccotti, G., Hynes, J.T. and Kapral, R. Constrained Reaction Coordinate Dynamics for the Simulation of Rare Events. Chem Phys Lett 1989;156:472–477. 10.1016/S0009-2614(89)87314-2

73. Sprik, M. and Ciccotti, G. Free energy from constrained molecular dynamics. J Chem Phys 1998;109:7737–7744. 10.1063/1.477419

74. Maragliano, L., Ferrario, M. and Ciccotti, G. Effective binding force calculation in dimeric proteins. Molecular Simulation 2004;30:807–816. 10.1080/0892702042000270205

75. Hirakawa, T., Bowler, D.R., Miyazaki, T., Morikawa, Y. and Truflandier, L.A. Blue moon ensemble simulation of aquation free energy profiles applied to mono and bifunctional platinum anticancer drugs. J Comput Chem 2020;41:1973–1984. 10.1002/jcc.26367

76. Brüssel, M., di Dio, P.J., Muñiz, K. and Kirchner, B. Comparison of Free Energy Surfaces Calculations from Molecular Dynamic Simulations at the Example of Two Transition Metal Catalyzed Reactions. International Journal of Molecular Sciences 2011;12:1389–1409. 10.3390/ijms12021389

77. Borisek, J. and Magistrato, A. All-Atom Simulations Decrypt the Molecular Terms of RNA Catalysis in the Exon-Ligation Step of the Spliceosome. Acs Catalysis 2020;10:5328–5334. 10.1021/acscatal.0c00390

78. Casalino, L., Nierzwicki, L., Jinek, M. and Palermo, G. Catalytic Mechanism of Non-Target DNA Cleavage in CRISPR-Cas9 Revealed by Molecular Dynamics. Acs Catalysis 2020;10:13596–13605. 10.1021/acscatal.0c03566

79. De Vivo, M., Dal Peraro, M. and Klein, M.L. Phosphodiester cleavage in ribonuclease H occurs via an associative two-metal-aided catalytic mechanism. Journal of the American Chemical Society 2008;130:10955–10962. 10.1021/ja8005786

80. Humphrey, W., Dalke, A. and Schulten, K. VMD: Visual molecular dynamics. Journal of Molecular Graphics & Modelling 1996;14:33–38. 10.1016/0263-7855(96)00018-5

81. Vosko, S.H., Wilk, L. and Nusair, M. Accurate Spin-Dependent Electron Liquid Correlation Energies for Local Spin-Density Calculations - a Critical Analysis. Can J Phys 1980;58:1200–1211. 10.1139/p80-159

82. Stephens, P.J., Devlin, F.J., Chabalowski, C.F. and Frisch, M.J. Ab-Initio Calculation of Vibrational Absorption and Circular-Dichroism Spectra Using Density-Functional Force-Fields. J Phys Chem 1994;98:11623–11627. 10.1021/j100096a001

83. Zhao, Y. and Truhlar, D.G. The M06 suite of density functionals for main group thermochemistry, thermochemical kinetics, noncovalent interactions, excited states, and transition elements: two new functionals and systematic testing of four M06-class functionals and 12 other functionals. Theor Chem Acc 2008;120:215–241. 10.1007/s00214-007-0310-x

84. Zhao, Y. and Truhlar, D.G. A new local density functional for main-group thermochemistry, transition metal bonding, thermochemical kinetics, and noncovalent interactions. J Chem Phys 2006;125. 10.1063/1.2370993

85. Frish, M.J. and al, e. Gaussian 09, Revision C.01. Gaussian, Inc., Wallingford CT. 2010.

86. Yu, M. and Trinkle, D.R. Accurate and efficient algorithm for Bader charge integration. J Chem Phys 2011;134:064111. 10.1063/1.3553716

87. Zhang, W., Qu, J., Liu, G.H. and Belmonte, J.C.I. The ageing epigenome and its rejuvenation. Nat Rev Mol Cell Biol 2020;21:137–150. 10.1038/s41580-019-0204-5

88. Zabransky, D.J., Jaffee, E.M. and Weeraratna, A.T. Shared genetic and epigenetic changes link aging and cancer. Trends Cell Biol 2022;32:338–350. 10.1016/j.tcb.2022.01.004

89. Dubey, S.K., Dubey, R. and Kleinman, M.E. Unraveling Histone Loss in Aging and Senescence. Cells 2024;13:320. 10.3390/cells13040320

90. Batchelor, L.K., De Falco, L., Dyson, P.J. and Davey, C.A. Viral peptide conjugates for metal-warhead delivery to chromatin. RSC Adv 2024;14:8718–8725. 10.1039/d4ra01617c

